# Conserved signaling pathways antagonize and synergize with co-opted *doublesex* to control development of novel mimetic butterfly wing patterns

**DOI:** 10.1101/2022.09.20.508752

**Authors:** Nicholas W. VanKuren, Meredith M. Doellman, Sofia I. Sheikh, Daniela H. Palmer Droguett, Darli Massardo, Marcus R. Kronforst

## Abstract

Novel phenotypes are increasingly recognized to have evolved by co-option of conserved genes into new developmental contexts, yet the impact of co-option on existing developmental programs remains obscure. Here we provide insight into this process by characterizing the consequences of *doublesex* co-option on wing color pattern development in *Papilio* swallowtail butterflies*. doublesex* is the master regulator of insect sex differentiation but has been co-opted to control the switch between discrete mimetic and non-mimetic, male-like color patterns in *Papilio polytes* and its close relatives. Here we show that development of the mimetic color pattern in *P. polytes* is caused by a pulse of *dsx* expression early in female wing development that results in a corresponding pulse of differential expression that both alters color pattern development and quickly becomes decoupled from *dsx* expression itself. Differentially expressed genes were enriched in canonical Wnt and Hedgehog signaling pathway genes, but case studies of key genes using RNAi and antibody stains suggested opposing, novel roles for the two pathways in mimetic color pattern development. The pulse of Dsx expression caused Engrailed, the key transcription factor effector of Hh signaling, to gain anterior expression in early pupal wing development. However, Dsx and En became decoupled by mid-pupal development when En pre-figured melanic and red patterns and Dsx pre-figured white patterns. In contrast, Wnt signaling antagonizes Dsx in restricted regions of the wing to refine the mimetic color pattern. Our results therefore provide strong experimental evidence that *dsx* co-option significantly altered spatiotemporal activities of conserved wing patterning pathways to promote and refine the development of a novel adaptive color pattern. Altogether, our findings provide strong evidence for how co-opted genes can both cause and elicit changes to established gene regulatory networks during the evolution and development of novel phenotypes.

## INTRODUCTION

It is now well appreciated that evolutionary novelties frequently arise from co-option of existing genes and gene regulatory networks (GRNs) into novel contexts, yet the molecular mechanisms underlying this process remain obscure [1–3]. Co-option has been particularly well documented to drive the evolution of novel morphological adaptations, from vertebrate lens crystallins [4] to stamen morphology in angiosperms [5] and tetrapod digits and feathers [6]. Current models describe gene co-option as a process in which a *cis*-regulatory change confers a novel expression pattern on a regulatory gene that results in activation of the gene and its downstream targets in a novel context, producing a developmental program and phenotype that can be refined by natural selection. While numerous examples of co-option caused by *cis*-regulatory changes have now been identified, the immediate and long-term effects of co-option on existing developmental programs and GRNs remain essentially unknown [7].

Classic examples of gene co-option come from studies of the evolution and development of butterfly wing color patterns, where patterns such as eyespots, bands, and rays evolved by the spatial and temporal redeployment of deeply conserved regulatory genes [8–11]. These genes, including multiple components of ancient animal signaling pathways such as Wnt, Hedgehog, and Notch, continue to perform their ancestral functions specifying the major axes of the wing, but have been repeatedly co-opted into developmental programs that function early in wing development to specify novel color patterns [12–17]. What has not yet been made clear is how novel expression patterns of these co-opted genes cause changes to existing color pattern networks to produce these new phenotypes.

Here we investigate the genes and GRNs controlling wing color pattern development in the swallowtail butterfly *Papilio polytes. Papilio polytes* and its close relatives have co-opted *dsx* from its role as the master regulator of insect sex differentiation to control the switch between discrete female wing color patterns [18,19]. While male *P. polytes* develop a single non-mimetic color pattern, females develop either a male-like pattern or a novel mimetic pattern, and the switch between female color patterns is completely controlled by a novel dominant *dsx* allele (Fig 1A) [20,21]. Recent work has identified additional candidate genes that function downstream of *dsx* to help specify the mimetic pattern using developmental stage-specific RNA-seq and RNAi. Iijima et al. [22] showed that the Wnt signaling ligands *wingless* and *Wnt6* function downstream of *dsx* to help specify mimetic red and white patterns, implicating this pathway in mimetic color pattern specification. Komata et al. [23] further showed that genes physically near to *dsx* in the genome, including *sirtuin-2* and *UXT*, help to refine the mimetic color pattern. But while these RNAi experiments clearly demonstrate that mimetic pattern development depends on these candidate genes, the molecular mechanisms by which *dsx* alters spatiotemporal activities of those genes and the development GRNs they comprise remain unknown [22,24].

**Figure 1.**
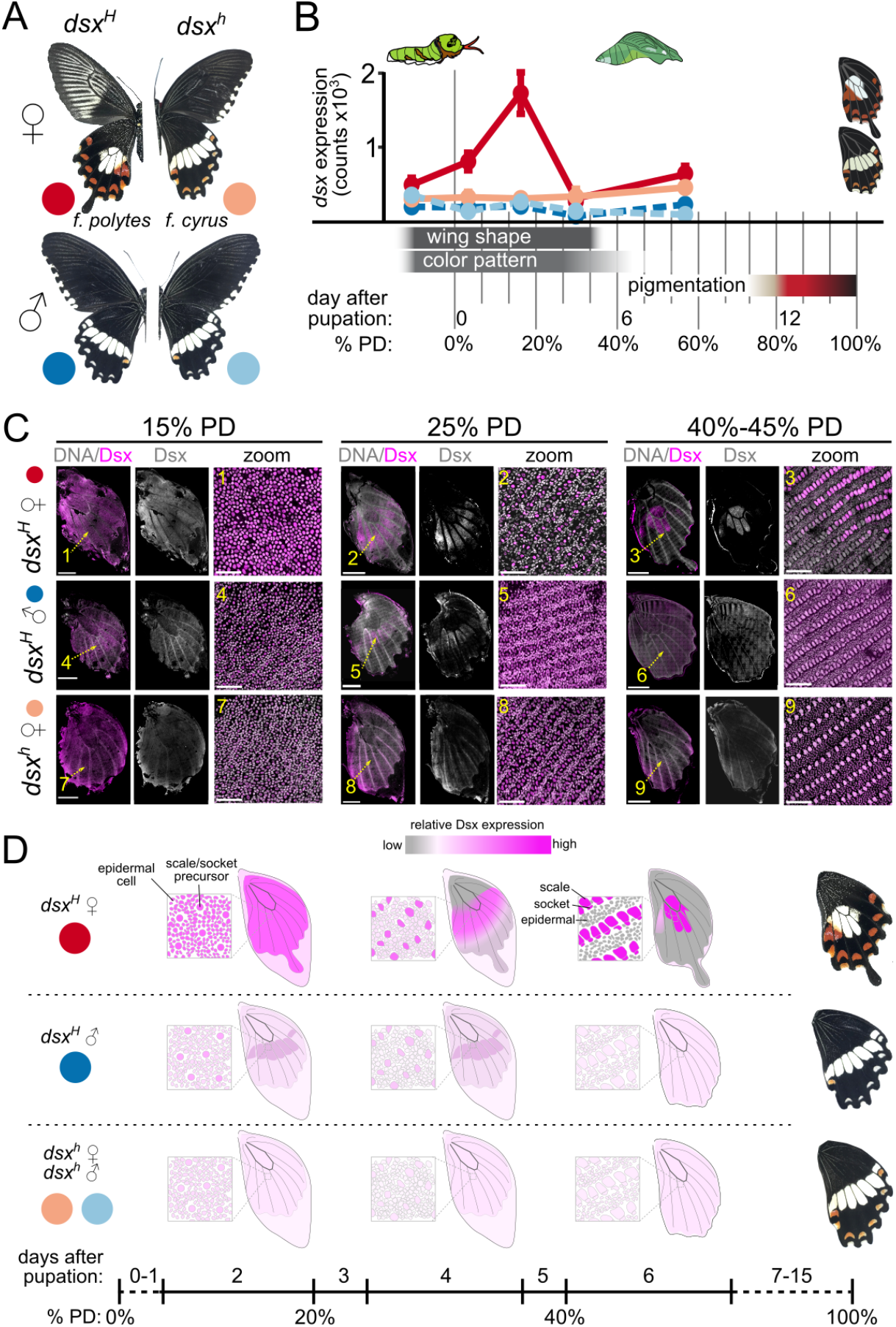
Dynamic *doublesex* expression patterns across *Papilio polytes* hindwing development. A) *Papilio polytes* color patterns and *dsx* genotypes. *dsx^H^* and *dsx^h^* are the mimetic and non-mimetic alleles, respectively. B) *dsx* expression in bulk hindwing RNA-seq data in relation to phases of hindwing development (see Fig 2). All four major *dsx* isoforms were expressed at each stage in all four groups, but the spike of *dsx* expression was primarily driven by female isoform 2 (Fig S2). C) Anti-Dsx antibody staining at select stages of hindwing development. Scale bars: 2 mm for full wings, 50 μm in zooms. D) Schematic of select stages of wing development and observed Dsx expression patterns. Scale cells produce the mature scales that comprise the adult wing color pattern; socket cells secure scale cells in the wing. These two cells derive from division of a single precursor cell around 40% PD. Epidermal cells comprise the bulk of the wing structure but do not contribute to color pattern. Additional stains can be found in Fig S1.

Here we provide insight into the consequences of gene co-option by reconstructing wing development GRNs, testing the effects of key genes on color pattern development using RNAi, and characterizing spatial patterns of Dsx expression across wing development using antibody stains. We use comparisons between sexes and individuals homozygous for different *dsx* alleles to disentangle the ancestral and co-opted roles of *dsx* at multiple layers of the color pattern development program. We identify temporal and spatial expression pattern differences between *dsx* alleles, identify differential gene expression and GRN differences associated with the mimicry switch, experimentally test the effects of altered signaling pathways on color pattern development, and perform a detailed study of one key effector of the mimicry switch, the transcription factor *engrailed*. Altogether, our results show that differential *dsx* expression alters the strength and patterning of Wnt and Hedgehog signaling between mimetic and non-mimetic butterflies and that these pathways have evolved to antagonize and cooperate with *dsx*, respectively, to produce novel mimetic color patterns.

## RESULTS

### Dynamic spatiotemporal expression patterns of Dsx

We first aimed to fully characterize the dynamics of Dsx expression in developing wings to understand where and when Dsx functions during color pattern development. All of the experiments in this paper used male or female butterflies that were homozygous for the nonmimetic or mimetic *dsx* alleles, allowing us to assay expression of each *dsx* allele independently and to disentangle the effects of sex and genotype on gene expression (Fig 1A). We first assayed Dsx expression using antibody stains in hindwings at five developmental stages in the four groups of butterflies. Dsx staining was uniform across the hindwing in all butterflies at 15% of pupal development (PD), with stronger overall staining in mimetic females consistent with RNA-seq data and previous qRT-PCR results (Fig 1) [22,23]. Non-mimetic Dsx remained weak and uniform across wings in males and females throughout development (Fig 1C; Fig S1). However, mimetic Dsx exhibited dynamic expression patterns in both sexes. By 40% PD in females, mimetic Dsx was restricted to scale and socket cell nuclei in regions of the wing that become white in the adult butterfly, with weaker staining in epidermal cell nuclei in some medial regions that become predominantly red. Stains at intervening days showed a smooth transition between these two general patterns (Fig 1; Fig S1). Mimetic Dsx staining was weak in males from 15% −40% PD, but was noticeably enriched in scale/socket precursor cell nuclei in regions that become pale yellow (Fig 1; Fig S1).

Thus, the mimetic *dsx* allele has gained novel expression in scale cells and their precursors in regions that become white and pale yellow in adults. Notably, mimetic Dsx expression did not fully pre-figure the mimetic color pattern at any of the developmental stages that we investigated, suggesting that co-opted Dsx expression in mimetic female wings alters color pattern development programs that continue to function long after Dsx expression is lost.

### An early spike of differential expression alters trajectories of conserved signaling pathway genes

Dsx stains suggested that differences in spatial expression patterns of the two alleles do not fully explain the color pattern switch. We next sought to identify genes and GRNs downstream of Dsx that may help effect the mimicry switch by generating and analyzing RNA-seq data from five developmental stages spanning the major phases of color pattern specification (Fig 1B; Table S1; Figs S3, S4). Consistent with our staining results, RNA-seq showed that *dsx* is low- to moderately expressed in male and non-mimetic female hindwings across early- to mid-pupal development, but that the mimetic allele undergoes a unique pulse of expression in mimetic females at 15% PD (Fig 1, ref. [19]). All four *dsx* isoforms are expressed in each group, but the spike of *dsx* expression in mimetic females is primarily driven by female isoform 2 (Fig S2).

We identified genes differentially expressed (DE) between mimetic and non-mimetic butterflies using two complementary approaches that disentangle the effects of sex and *dsx* genotype on gene expression (Fig 2; Table S2; Methods). First, we found 904 genes DE at one or more developmental stages by analyzing each stage separately, with 98.2% of these stage-specific DE genes coinciding with the pulse of mimetic *dsx* expression at 15% PD (Fig 2A). The majority of stage-specific DE genes (65.9%) were down-regulated in mimetic females relative to non-mimetic females (Fig 2A). We also identified 1598 genes with significantly different temporal expression profiles in mimetic females relative to non-mimetic females (878) or all nonmimetic butterflies (720; Fig 2B). It is important to note that both temporal contrasts (mimetic female to non-mimetic female and mimetic female to all) are necessary for identifying genes involved in the mimicry switch: the mimetic female-all comparison identifies genes that are DE specifically in mimetic females, while the female-female comparison identifies genes that are DE specifically in non-mimetic females. While superficially similar, there are important differences in scale color and structure between non-mimetic females and males that suggest their developmental programs may also differ. Stage-specific and temporal analysis results were largely non-overlapping, with stage-specific tests identifying genes with a single sharp peak or trough in expression and temporal tests identifying genes with subtler expression profile differences (Fig 2C, D). Altogether, these results suggested that the mimicry switch depends on an early spike of *dsx* expression that has both acute and long-term effects on gene expression in developing female hindwings.

**Figure 2.**
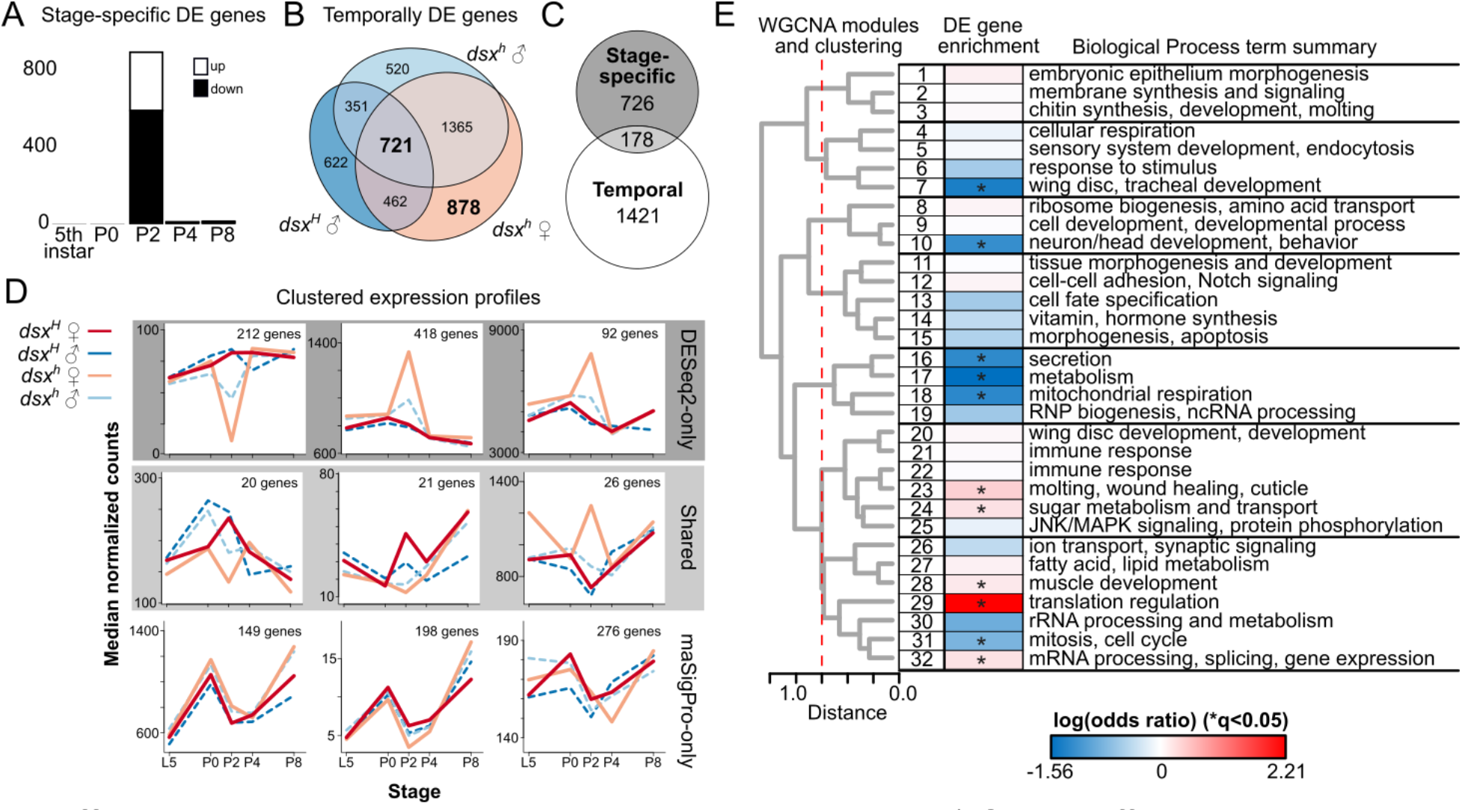
Differential expression underlying the mimicry switch. A) Genes differentially expressed between mimetic and non-mimetic females at each stage, identified using DESeq2 (overall FDR < 0.01). Up/down is mimetic relative to non-mimetic females. B) Euler diagram of genes with significantly different expression profiles in mimetic *dsx^H^* females relative to the indicated groups, identified using maSigPro (overall FDR < 0.01). C) Euler diagram of overlap between DESeq2 and maSigPro results. D) Median expression profiles of three largest clusters for each set in C. E) Relationships, DE gene enrichment, and Gene Ontology BP term enrichment of co-expressed gene modules identified using WGCNA (full results in Fig S5 and Table S3). Module numbers are arbitrary. Dashed red line indicates cuts to define metamodules. *Benjamini-Hochberg corrected Fisher Exact Test *p-*value < 0.05.

We next aimed to identify the developmental GRNs and pathways that were affected by the early pulse of *dsx* expression. We reconstructed the hindwing development co-expression network using WGCNA, then identified sets of co-expressed genes (modules) enriched with DE genes (Fig 2E; Fig S5; Table S3) [25,26]. Five of the 32 identified modules were significantly enriched with DE genes (Fig 2E; Table S3). Consistent with the idea that the pulse of *dsx* expression has wide-ranging effects on color pattern development programs, the two most significantly enriched modules (29 and 32) are significantly enriched with Gene Ontology terms for gene expression regulation at the levels of both transcription and translation (Fig 2E; Table S3).

Conserved signaling pathways such as Wnt, Hedgehog (Hh), *TGF-β*, and Notch have been repeatedly co-opted into wing color patterning networks from their ancestral roles specifying tissue and cellular polarity [8–15,17]. In addition, previous RNAi work in *P. polytes* clearly showed that the Wnt signaling ligands *wg* and *Wnt6* play a role in mimetic color pattern development [22]. We were therefore interested to know the extent to which genes in these pathways were involved in the Dsx mimicry switch, and how *dsx* co-option had altered their activities. DE genes were indeed significantly enriched with canonical Wnt (*χ*^2^ = 8.51, 1 d.f.; *p* = 0.0035) and Hh (*χ*^2^ = 9.77, 1 d.f.; *p* = 0.0018) pathway components, including multiple Wnt ligands, core components of the *β-catenin* / *cubitus interruptus* regulator complex, and key transcription factor effectors (Table 1; Tables S4-S6). These results altogether suggest that mimetic Dsx causes a fundamental shift in the strength or pattern of these core signaling pathways across the developing mimetic wing.

**Table 1.**
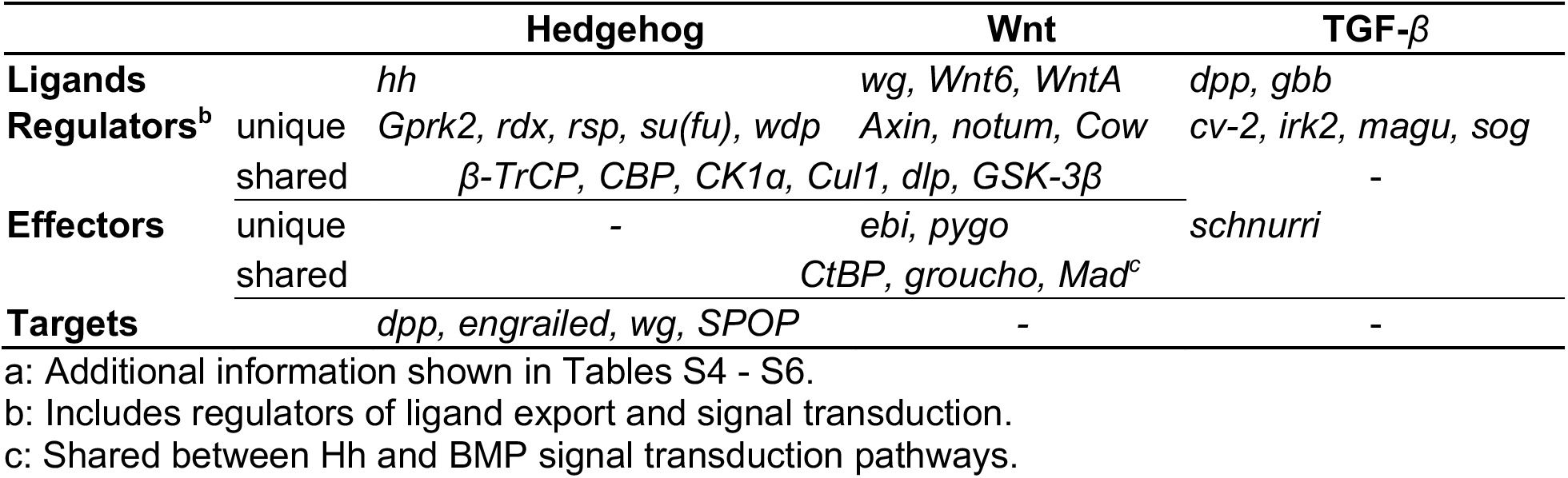
Signaling pathway components DE between mimetic and non-mimetic females^a^.

### Canonical Wnt signaling antagonizes mimetic color pattern development

We next tested whether differential expression of these signaling pathway components caused color pattern differences using RNAi. siRNAs were injected into one hindwing at pupation, then pupae were allowed to finish development and emerge as adults, following ref. [27]. This method yields strong, long-lasting, and widespread knockdowns of target genes (Fig S6). Consistent with previous studies, *dsx* RNAi in mimetic females resulted in a complete switch from the mimetic to the non-mimetic color pattern, but no effect on non-mimetic color pattern development in either sex (Fig 3A) [19]. Results were similar using siRNAs targeting either exon 3 (all female isoforms) or exon 5 (all isoforms except female isoform 2C; Fig S2; Dataset S1).

**Figure 3.**
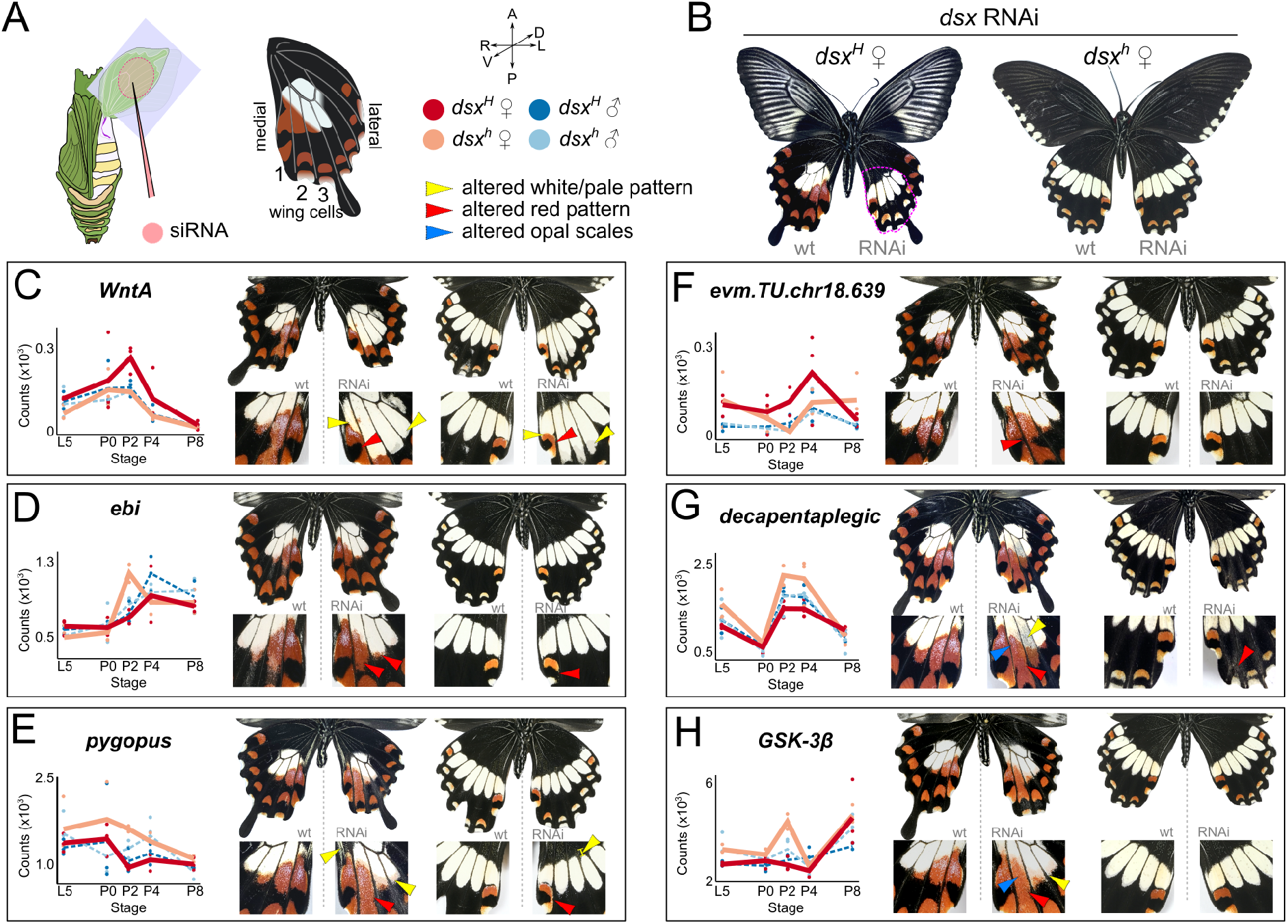
RNAi shows DE genes primarily affect mimetic color pattern development. A) siRNAs are injected and electroporated into the ventral left hindwing at pupation. Landmarks, orientations, and keys applicable to all panels. B) *dsx* RNAi phenotypes in mimetic and nonmimetic females. The area typically affected by RNAi injections is circled in the mimetic female. C-H) Expression patterns from bulk RNA-seq data and phenotypes in mimetic and non-mimetic females for six target genes. Note that bulk RNA-seq data cannot distinguish between changes in gene expression level and pattern. All images are of the ventral surface and all comparisons should be made between color patterns on the right wing (wild-type, wt) and left wing (RNAi) of the same individual. Full phenotypes and over 70 additional RNAi individuals, including males of both genotypes, are found in Dataset S1.

Previous work established that *wg* and *Wnt6* are required for development of submarginal red patterns and some proximal white patterns in mimetic females [22]. We next tested the effects of a set of DE genes from within the Wnt and Hh signaling cascades to gain insight into the effects of *dsx* on these conserved pathways and to determine their roles in color pattern development in general. We therefore used RNAi in the four groups of butterflies to knock down the signaling ligands *WntA* and *decapentaplegic;* the shared regulator *GSK-3β;* transcription factor effectors *ebi, pygopus, engrailed*, and *invected;* and one DE gene with unknown function, *evm.TU.chr18.639* (Fig 3).

RNAi of canonical Wnt components resulted in a striking expansion of red patterns and opal scales, which are typically a rare scale type co-localized with red patches, in mimetic female wings (Fig 3). While *ebi* RNAi resulted in mild increases of red scales and decreases in opal scales in the medial two wing cells, *pygopus* and *GSK-3β* RNAi caused most melanic scales in the second wing cell to become red; extensive loss of opal scales; and extension of the white patch over central red patterns (Fig 3). *evm.TU.chr18.639* RNAi caused phenotypes similar to *ebi* knockdowns, suggesting that it may also function in Wnt signal transduction. Importantly, we observed mild or no effects of RNAi of these genes in non-mimetic butterflies, regardless of their *dsx* genotype, suggesting that mimetic Dsx has caused the evolution of a mimetic female-specific pattern of Wnt signaling that functions to restrict medial red and opal patterns promoted by the mimetic Dsx itself (Fig 3; Dataset S1). This conclusion is supported by 1) few, weak phenotypes from RNAi in non-mimetic butterflies and 2) the fact that RNAi never fully recapitulated the non-mimetic color pattern, except for *dsx* RNAi. If genes are active in these same regions in non-mimetic butterflies, then they are not altering default melanic color patterns. For example, *ebi* and *pygo* are positive regulators specific to the canonical Wnt signaling pathway but are overall downregulated in mimetic females, suggesting that these genes are spatially restricted in mimetic females. Uniquely, *WntA* RNAi caused massive distal expansion of white/pale patches in all butterflies, suggesting it has a more general role establishing the distal boundaries of those patterns, similar to its function in some Nymphalidae [28] (Fig 3; Dataset S1).

RNAi targeting genes outside of the canonical Wnt pathway affected both red and white patterns. RNAi of *decapentaplegic*, a key target of Hh signaling and one of three TGF-*β* ligands, yielded a phenotype similar to *GSK-3β* knockdowns, but with stronger effects near the wing margin. *dpp* RNAi also affected margin patterns in non-mimetic butterflies (Fig 3G). In general, we observed few alterations to marginal patterns, but antibody stains in RNAi wings showed this may be due RNAi mostly affecting the middle 50%-75% of the wing (Fig 4, Figs S6, S7). Finally, RNAi of Hh transcription factor effectors *engrailed* and *invected* had fundamentally different effects on color pattern than the genes discussed so far and we investigated them further below.

**Figure 4.**
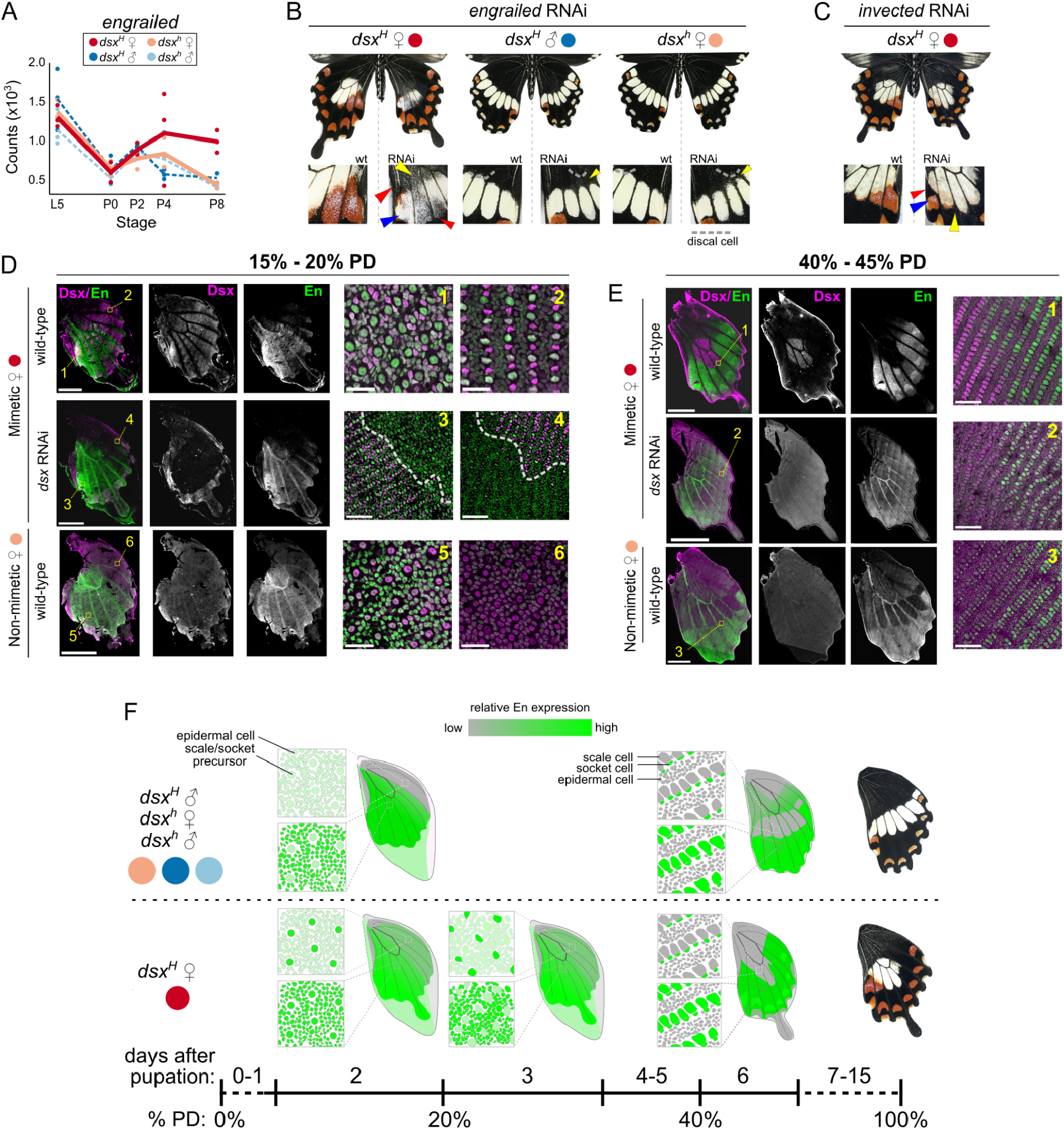
The Doublesex mimicry switch acts through Engrailed. A) Temporal *en* expression patterns. B) *en* RNAi phenotypes. C) *inv* RNAi phenotypes. D) En and Dsx antibody staining patterns at 15-20% PD. Dashed lines in zoom images mark the edges of *dsx* RNAi clones. E) En and Dsx antibody staining patterns at 40%-45% PD. Brightness levels between pairs of zoom images in D and E are comparable because they were taken with the same settings and are not adjusted. Scale bars: 2 mm for full wings; 25 um for D1, 2, 5, 6; and 50 um for remaining for zooms. F) Schematic of En expression across early- to mid-pupal hindwing development. See additional stains in Fig S6.

Altogether, RNAi from this work and others supports a central role of Wnt signaling in antagonizing the effects of the novel mimetic *dsx* allele [22]. We add to this previous work by studying the effects of knocking down key regulators and effectors. In contrast to *ebi* and *pygo, GSK-3β* is a negative regulator of both Wnt and Hh signaling, making it difficult to disentangle the effects of *GSK-3β* RNAi on each pathway. In general, there is an enormous amount of cross-talk between the Wnt and Hh pathways [29], from shared regulators such as *GSK-3β* and *CBP* to co-dependent expression of *wg* and *hh* themselves, that could also allow some degree of compensation. What is clear is that these genes and pathways are critical for the proper development of a good mimetic color pattern.

### The Dsx mimicry switch acts through Engrailed

In contrast to RNAi of canonical Wnt components, RNAi of *engrailed* and *invected* caused altered central mimetic patterns that superficially resembled the non-mimetic color pattern (Fig 4A). *engrailed* RNAi had strong effects in all butterflies (Fig 4B). In non-mimetic butterflies, *engrailed* RNAi caused the pale band to shift distally, suggesting that *engrailed* normally specifies the proximal-distal positioning of these windows in the melanic background (Fig 4B). The shape of the pale patches remained mostly unchanged, but the shift could be so drastic that marginal and submarginal patterns merged (Dataset S1). In contrast, *en* RNAi in mimetic females caused complete loss of central red patches and restriction of the central white patch to a band reminiscent of the non-mimetic color pattern (Fig 4B). However, this restricted patch comprised primarily opal and gray scales instead of the typical white or pale yellow scales. These gray scales are normally found at the boundaries of white/pale and melanic regions in wild-type wings. The medial red patch now comprised mainly opal scales. Altogether, *en* RNAi strongly suggests this gene, the key transcription factor effector of the Hh signaling pathway, gained a novel role in mimetic color pattern development that helps specify the location of novel color pattern elements and alters scale colors within those elements.

Consistent with this idea, we found striking differences in En antibody staining patterns between mimetic and non-mimetic pupal wings. En was expressed in all nuclei of the posterior two-thirds of late larval instar wing discs in all butterflies, consistent with its ancestral role specifying posterior compartments of appendages [17,30] (Fig S7). Posterior compartment expression was maintained into early pupal development in all butterflies, where En marked epidermal cells (20% PD; Fig 4C). However, En was uniquely expressed in alternating scale/socket precursors across the distal half of the mimetic female wing. Dsx was expressed in all scale/socket precursors in this region, suggesting that Dsx and additional cell non-autonomous signals coordinate this novel En expression domain. *dsx* RNAi caused complete loss of the novel En expression pattern, but did not alter En expression in epidermal cells, showing that the pulse of mimetic *dsx* expression at 15% PD is necessary for En expression in scale/socket precursors (Fig 4). Males carrying the mimetic *dsx* allele also expressed En in some scale/socket precursors in the distal half of the wing, excluding future pale patches, suggesting mimetic *dsx* is not completely impotent in this sex (Fig S7).

The pulse of mimetic *dsx* expression had long-term effects on *en* expression patterns. By 40%-45% PD in non-mimetic butterflies, En expression expanded to scale and socket cells across the wing, with much weaker expression in future pale yellow patches (Fig 4D). En expression was strongest in the distal half of the wing, and slightly higher in future red patches, supporting the *en* RNAi data and the hypothesis that En specifies the proximal-distal positioning of the pale patches in non-mimetic butterflies. En was also expressed in scale cells across the distal half of the wing in mimetic females at 40%-45% PD, but was highly enriched in all future red patches and excluded from Dsx-positive scale cells in the white patch (Fig 4D). *dsx* RNAi in mimetic females resulted in a non-mimetic-like staining pattern where En was only weakly expressed in future pale patches. Thus, by mid-pupal stages En and Dsx expression patterns are decoupled. *en* RNAi did not affect Dsx expression patterns at 20% or 40% PD (Fig S8). These results provide strong support for our hypothesis that the spike of mimetic *dsx* expression and DE genes at 15% PD has long-lasting effects on the strength and pattern of gene expression in the developing wing that become independent of *dsx* expression itself (Fig 2).

*engrailed* and *invected* are frequently investigated together because these paralogs are co-regulated in *Drosophila* and loss-of-function of one paralog can be at least partially compensated for by the other. However, the two genes share only 50% coding sequence identity, 19% protein sequence identity, and differentially function in development of numerous fly tissues. We were therefore interested to know if *inv* had also been co-opted into the *dsx* switch. *Papilio polytes engrailed* was DE in mimetic females relative to all non-mimetic butterflies, remaining highly expressed in later developmental stages (Fig 4A). *inv* was not DE, but was slightly elevated in mimetic females at the same stages as *en*. *inv* RNAi also altered the mimetic color pattern, producing a non-mimetic-like pattern in wing cell 1, distal shift of the white patch, and reduced submarginal red patches. Unlike *en* RNAi, we observed no increase in opal or gray scale frequencies. *en* and *inv* therefore both participate in the mimicry switch but to different degrees. It is unlikely the *en* and *inv* reagents we used cross-reacted because of their high sequence divergence, different RNAi phenotypes, and the observed complete loss of anti-En staining in *en* RNAi wings (Fig S8). *en* and *inv* RNAi and staining patterns altogether suggested these ancient paralogs cooperate with mimetic *dsx* to produce the novel color pattern.

## DISCUSSION

Co-opted genes are expected to have immediate pleiotropic effects on local GRNs that must be mitigated for the co-opted allele to be maintained and fixed in the population. However, those effects and the long-term consequences of co-option on the structures and functions of local GRNs remain poorly characterized [7]. Here we provide insight into the consequences of co-option through comparative, functional, and experimental analyses of the *dsx-*mediated mimicry switch in *P. polytes*.

The genetic evolution of the *dsx* mimicry switch is through to have occurred through a series of steps, including: 1) the accumulation of mutation(s) that conferred a novel expression pattern in developing wings on *dsx*; 2) an inversion that locked the causative mutation(s) into a single mimetic haplotype; and 3) additional mutation accumulation in the mimetic allele that refined its function in mimetic color pattern development while maintaining its ancestral role in sex differentiation. Our results and others suggest that the ancestral, non-mimetic *dsx* allele plays a limited role in color pattern development: it is lowly expressed across the developing wing and *dsx* RNAi has minimal effects on color pattern (Dataset S1) [19,23,31]. Thus, the mimetic allele has gained a novel, dynamic expression pattern in the developing hindwing. While our RNA-seq data show that pulse of mimetic *dsx* expression is driven by female isoform 2 (Fig S2), it is not yet clear if alternative splicing of *dsx* plays a role in the mimicry switch like it does in sexual differentiation. Results suggesting that female isoform 3 is the key player in the mimicry switch relied on an siRNA that perfectly matches all three female isoforms [23]. Furthermore, there is no evidence that different Dsx isoforms (male and female) bind to different target sequences and in fact have identical DNA binding domains. In fact, work in *Drosophila* has clearly demonstrated that Dsx binds to the same sites in both sexes and different tissues, and that it is the local regulatory environment and binding partners that dictate the effects of Dsx binding on nearby gene expression [32].

The most significant difference between *dsx* alleles is the pulse of mimetic *dsx* expression early in pupal development, which causes a corresponding spike in genes that are differentially expressed between mimetic and non-mimetic females (Fig 1B, Fig 2A). However, our developmental time series showed that the majority of the DE genes had altered temporal expression profiles, strongly suggesting that the early pulse of co-opted *dsx* expression fundamentally alters regulatory gene expression early in development that cascades to later stages. This is supported by network and GO analyses (Fig 2) that show gene modules associated with transcription and translation regulation are enriched with genes that are DE in mimetic females. Interestingly, Dsx and En stains suggest that GRN activities at later stages are decoupled from *dsx* itself, showing that even a brief period of co-opted gene activity can significantly alter local regulatory environments. While we could not properly compare our results with previous studies because of selective presentation [22,23] or different experimental designs [22–24], we found 32% of genes presented by Iijima et al. [22] were also found DE here. Our DE results do not rely on RNAi, and bulk RNA-seq has low power to detect spatially restricted genes, so this is not necessarily surprising. Application of spatial transcriptomics and functional genomics approaches will be critical for assaying *dsx* function and regulation across the developing wing.

Several molecular mechanisms appear to have evolved to limit the mimetic allele’s function since it formed about 1.7 mya [21], including the gain or loss of *cis-*regulatory elements that control the early pulse and/or spatially restricted expression patterns and alterations of the strength or pattern of canonical Wnt signaling. Our observations that Wnt component RNAi rarely altered non-mimetic color pattern development in males or females strongly supports the idea that altered Wnt signaling is a direct result of *dsx* co-option. Interestingly, males also express mimetic *dsx* and exhibit slightly altered *en* expression, suggesting that mimetic *dsx* expression is not fully refined, or cannot be fully refined due to pleiotropy. The *cis*-regulatory differences between the two *dsx* alleles are currently unknown, but their identification will provide crucial insight into the process by which the mimetic *dsx* allele gained its novel expression pattern and how it was refined by subsequent modifications to the color pattern GRN.

Our findings suggest a process for mimetic color pattern development that depends on partial suppression of Dsx function by Wnt signaling: a pulse of *dsx* early in development triggers activation of a mimetic color pattern development program whose spatial activity is refined by Wnt and Hh signaling despite being uncoupled from Dsx expression itself. This resembles a recent model put forward by Komata et al. [23], who showed that genes near the *dsx* inversion, especially *UXT* and *sir2*, antagonize mimetic Dsx function. However, the similarity between phenotypes caused by *sir2* RNAi and our Wnt pathway RNAi, and observations that *sir2* directly interacts with *β-catenin* in vertebrates [33], may suggest that *sir2* affects color pattern through its effects on Wnt signaling. In addition, it is not clear if these genes affect nonmimetic color pattern development. These results more generally highlight that we do not know 1) the genes that Dsx directly regulates or 2) the directionality of the interactions between the genes in the *dsx*-orchestrated gene network, critical information for comparing color pattern development before and after *dsx* co-option. Future work identifying Dsx binding sites and using double RNAi knockdowns will be necessary to establish directionality in the mimetic network.

Finally, the Hedgehog transcription factor *engrailed* provides unique insight into the consequences and resolution of co-option. *engrailed*, and possibly *invected*, pre-figure eyespots and marginal patterns in some nymphalid larval wing discs [9,13,34], and were repeatedly coopted into dipteran color pattern networks [35,36]. Dufour et al. [36] showed that the interactions between genes in the Hh signaling pathway change over wing development, and *engrailed* in particular has been co-opted to produce novel wing color patterns in some Drosophilidae without affecting most other components of the Hh GRN. Interestingly, the novel En expression domain in *P. polytes* does not depend solely on *dsx* because the two genes are not perfectly coexpressed (Fig 4). Hh and Wnt negatively feed back on each other during segment polarity specification, and this may be another manifestation of that interaction. Importantly, En does not appear to be performing a fundamentally different function in non-mimetic and mimetic color pattern development. Instead, mimetic *dsx* alters the strength and pattern of En expression across the wing to shift the position of non-pigmented windows and the frequency of red scales. Notably, genetic variation near *engrailed* and *invected* has also been associated with complex female-limited mimicry polymorphism in *Papilio dardanus* [37,38]. Interestingly, the *en* RNAi phenotypes we observed are quite similar to the female-limited color pattern variation observed in *P. dardanus*, where distal expansion and shifts of central white patches distinguish the common morphs *cenea* and *hippocoonides* [38]. The *hippocoonides* allele is completely recessive, perhaps suggesting a lack of *en* expression and reflecting the *P. polytes en* RNAi phenotype.

## ACKNOWLEDGEMENTS

We thank Nipam Patel for providing antibodies, The University of Chicago Functional Genomics Facility (RRID:SCR_019196) and its staff for their help with sequencing, the University of Chicago greenhouse staff, and Ayşe Tenger-Trolander, Kelsey Stilson, Erick X. Bayala, and Adam Kuuspalu for technical assistance. This work was funded by NIH R35 GM131828 to MRK.

## AUTHOR CONTRIBUTIONS

NWV designed experiments, performed experiments, analyzed data, and wrote the manuscript. MMD, SIS, and DM performed experiments. DHPG designed experiments and performed experiments. MRK designed experiments and acquired funding.

## MATERIALS AND METHODS

### Butterfly care and pupa dissection

*Papilio polytes alphenor* pupae were purchased from Philippines breeders and grown in the University of Chicago greenhouses. Virgins were labeled on the hindwing with permanent marker, then genotyped for *dsx* alleles using a leg and custom TaqMan (Thermo Scientific) assays. Males and females were housed in separate 2 m^3^ mesh cages until setting up single-pair crosses between individuals homozygous for mimetic *dsx^H^* or non-mimetic *dsx^h^* alleles. *Citrus* shrubs were provided for oviposition and larval food. Pre-pupae were collected each morning, and photographed to monitor pupation times. Pupae were transferred to an incubator (70% RH, 25°C, 16h:8h light/dark cycle) within 8 hours after pupation (AP) to continue development. A pupa between 12 h and 24 h AP is a day 0 pupa (P0).

### Antibodies and staining

We raised a new antibody against *P. polytes* Dsx. We used the protein encoded by the first two exons because they comprise more than 80% of the full protein, are shared between all isoforms, and exhibit 94% identity between *dsx* alleles. Protein synthesis, conjugation, purification, and immunizations were all carried out by GenScript (USA). We confirmed the specificity of this antibody using RNAi and staining in pupal wing discs (Fig 4, Fig S6).

Pupal wings were dissected in room temperature PBS. Cuticle, forewings, and hindwings were dissected out as a single unit. After removing peripodial membranes covering hindwings, wings were fixed for 15 min in 3.7% formaldehyde in PBST (PBS + 0.1% Triton X-100). Wings were washed twice quickly, dissected out from the cuticle and transferred to 4-well plates, then washed 3 x 15 min in PBST. Wings were stored in PBST at 4°C until staining. Wings were blocked in 1% BSA in PBST for 1 hr at RT, then incubated in blocking buffer with primary antibody overnight at 4°C. Wings were washed twice quickly then 5 x 10 min in PBST at RT before adding secondary staining solution and incubating overnight at 4°C. Wings were washed twice quickly, 5 x 10 min, and 3 x 1 hr in PBST at RT, incubated in 50% glycerol/PBS for 30 min, then covered in Vectashield (Vector Laboratories) until mounting. Antibodies were used at 1:100 (4F11), 1:500 (α-Dsx), or 1:1000 (GαRb AlexaFluor-488, GαRb AlexaFluor-555, DαM AlexaFluor-555, DAPI). Samples were mounted in fresh Vectashield, using double-sided tape to make a coverslip bridge. Images were captured on a Zeiss LSM 710 confocal microscope at the University of Chicago Department of Organismal Biology & Anatomy, then processed using ImageJ. Whole wings were imaged using a 20X objective and a Z-stack / tile scan across the whole wing to capture the entire ventral surface. Images were then stitched and converted to maximum intensity projections in Zen or ImageJ. We took additional images with the 40X objective to characterize cellular localization. Images were assembled and adjusted for brightness and contrast in Inkscape.

### RNA-seq library construction and sequencing

Hindwings were dissected out in ice cold PBS, cleaned of peripodial membrane and cuticle, then immediately transferred to RNAlater (Ambion) before storage at −80°C until RNA extraction. One replicate comprised three hindwings from three individuals from a single cross. We extracted total RNA using TRIzol (Ambion), then depleted 18S, 5.8S, 28S, 12S, and 16S rRNAs from each sample using the RNase H method [39]. Depleted RNAs were used to construct sequencing libraries using the KAPA RNA HyperPrep Kit (KAPA Biosystems), amplified for 11 cycles, and sequenced 1×50 bp on an Illumina HiSeq 4000 at the University of Chicago Functional Genomics Facility (Table S1).

### Differential expression analysis

We assembled and annotated a *Papilio polytes alphenor* genome (see SI Materials and Methods), then quantified transcript expression levels using Salmon v1.4.0 [40] with bias correction and the full transcript annotation set, then imported and normalized quantification data using tximport 1.18.0 [41] and DESeq2 [42]. Differential expression analyses were performed using normalized gene-level quantification data. We identified genes at each developmental stage that respond to the mimetic *dsx* allele specifically in females using DESeq2. We used the following model for these tests, using *dsx^h^* males as the baseline (i.e. sex = 0, genotype = 0):

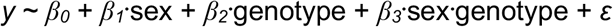

We identified genes with significant *β_2_*+*β_3_* terms, controlling FDR < 0.01 over all five stages. This is equivalent to defining four sex-genotype groups and performing the pairwise comparison between female groups at each stage.

We also identified genes with significantly different temporal expression profiles in mimetic females using *maSigPro* 1.64.0 with a quartic fit [43,44]. Significant genes were selected using a *q*-value cutoff of 0.01 for the `p.vector()` function, then variable selection performed using *p*-value < 0.05 in `T.fit()`. Finally, genes with good fits were defined as those with *R^2^* > 0.6. Genes were clustered by expression profile and using maSigPro functions that we modified.

### Co-expression network reconstruction and pathway enrichment

We reconstructed the hindwing development gene co-expression network using WGCNA and the gene-level quantification data from above. Adjacency and topological overlap matrices (TOMs) were constructed using signed Pearson coefficients. We performed GO enrichment analyses on each module using topGO v2.44.0 [45] and eggNOG GO assignments (see SI Materials and Methods), utilizing the Fisher Exact Test with adjusted *p*-value < 0.01 to identify enriched GO categories. We tested for enrichment of canonical Wnt and Hedgehog signaling pathway genes as defined by KEGG pathways 04310 and 04341, respectively. We identified all *alphenor* orthologs of the genes in those pathways using blastp, then performed FETs against all DEGs. The mappings and tests are shown in Tables S4 and S5. We tested whether modules were significantly enriched or deficient in switch genes using FETs.

### RNAi

RNAi experiments were performed as described previously by Ando and Fujiwara [27] with small modifications. We designed 24 - 27 nt long Dicer substrate siRNAs (DsiRNAs) using IDT’s DsiRNA design tool and the full transcript of the target gene, excluding any designs with off-targets (Table S7). We identified off-targets using primer-BLAST and defined them as any non-target transcripts with fewer than 5 mismatches to the DsiRNA. We injected 1.5 uL of 100 μM DsiRNA into the left hindwing near the discal cell and between veins Cu1 and M3, covered the injected area with PBS, and electroporated into the ventral epithelium using five 0.25 sec, 10V shocks spaced over 5 seconds. Pupae were then placed in an incubator in petri plates with moist paper towels to allow them to finish development or until dissection for stains. Full RNAi results can be found in Dataset S1.

