## Supplementary Information for "Conserved signaling pathways antagonize and synergize with co-opted *doublesex* to control development of novel mimetic butterfly wing patterns"

**SUPPORTING INFORMATION**

**Includes:**

Supplemental materials and methods

Figures S1 - S8

SI References

**Supplemental materials and methods**

*Papilio polytes alphenor genome assembly*

The published *P. polytes* reference genome was generated using *P. polytes polytes* from Japan, while our data is from *P. p. alphenor* from the Philippines. These two groups diverged ~1.7 million years ago and have ~5.1% nucleotide divergence (1), resulting in low mapping rates of our *alphenor* RNA-seq data to the published reference (2). We therefore generated a new, high-quality *alphenor* genome assembly based on the *P. polytes polytes* assembly and a recent chromosome-level *P. bianor* assembly.

We assembled a draft genome using PE100 sequencing from 29 individuals (PRJNA234541, excluding SRR1118138) and Platanus v2.0.2 (3, 4) (Table S8). Before assembly, we trimmed raw reads using TrimGalore! (<https://www.bioinformatics.babraham.ac.uk/projects/trim_galore/>), then removed reads containing over-represented sequences using FastQC (<http://www.bioinformatics.babraham.ac.uk/projects/fastqc/>) and an available python script (<https://github.com/harvardinformatics/TranscriptomeAssemblyTools/blob/master/RemoveFastqcOverrepSequenceReads.py>). We then assembled all processed reads using the default platanus2 pipeline. Next, we assigned raw *alphenor* scaffolds to the RefSeq *polytes* assembly (GCF_000836215.1) using RagTag v1.0.1 (5), then assigned these *alphenor* pseudoscaffolds and all other *alphenor* scaffolds that hit insect sequences in the NCBI nr database to *P. bianor* chromosomes (6).

Finally, we assembled *alphenor doublesex* alleles separately and substituted them into the final chromosome-level assembly. These additional steps were necessary because the individuals used for the full Platanus assembly were a mix of homo- and heterozygotes at the *dsx* locus - while the majority of the genome is homogeneous among these samples, the *dsx* alleles are very divergent except for the *dsx* coding region. We assembled all non-mimetic female samples (SRR1109784, SRR1111718, SRR1111828, SRR1111944, SRR1112070, SRR1112485, SRR1112619, SRR1117689, SRR1118114, and SRR1118137) using Platanus 2 and assigned scaffolds to the *polytes* assembly using RagTag. We then pulled out the *polytes* region corresponding to the *dsx* inversion, defined by Nishikawa et al. (2) as H_locus_nonmimetic_H_scaffold:1931762-2054949, and used it to replace the corresponding region in the chromosome-level *alphenor* assembly. Similarly, we assembled the mimetic *dsx* allele by assembling all mimetic female samples (SRR1118143-SRR1118152) using Platanus 2, assigning those scaffolds to the RefSeq assembly, and extracting the H_locus_mimetic_hetero scaffold. We added this scaffold as chrH in the final *alphenor* assembly.

We assembled the *alphenor* mitochondrial genome using NOVOplasty v4.2 (7) using sequencing data from SRR1108726 and the RefSeq mtDNA assembly for *polytes* (NC_024742.1) as the seed sequence. This resulted in a single circularized sequence of 15,247 bp.

*Transcriptome assembly and genome annotation*

We annotated the *alphenor* genome using EvidenceModeler 1.1.1 (8). We first assembled a high-quality transcript database using PASA (8), our SE50 data, and PE100 and SE50 data from Nallu et al. (9). After adapter trimming, we performed *de novo* and genome-guided assembly using Trinity v2.10.0 (10) and genome-guided assembly using StringTie v1.3.3 (11). RNA-seq data was also mapped to *alphenor* chromosomes using STAR 2.6.1d (12), and the resulting alignments used to generate genome-guided assemblies with Trinity and StringTie 1.3.1 (11). We combined *de novo* and genome-guided assemblies using PASA 2.4.1 (13). Evidence for protein-coding regions came from mapping the UniProt/Swiss-Prot (2020_06) database and all Papilionoidea proteins available in NCBI’s GenBank nr protein database (downloaded 6/2020) using exonerate (14). We identified high-quality multi-exon protein-coding PASA transcripts using TransDecoder (transdecoder.github.io), then used these models to train and run Genemark-ET 4 (15) and GlimmerHMM 3.0.4 (16). We also predicted gene models using Augustus 3.3.2 (17), the supplied *heliconius_melpomene1* parameter set, and hints derived from RNA-seq and protein mapping above. Augustus predictions with >90% of their length covered by hints were considered high-quality models. Transcript, protein, and *ab initio* data were integrated using EVM with the weights in Table S9.

Raw EVM models were then updated twice using PASA to add UTRs and identify alternative transcripts. Gene models derived from transposable element proteins were identified using BLASTp and removed from the annotation set. The final annotation comprises 17,342 genes encoding 26,991 protein-coding transcripts, containing 95% and missing 3% of endopterygota single-copy orthologs according to BUSCO v3 and OrthoDB v9 (Table S8). We functionally annotated protein models using eggNOG’s emapper-2.0.1b utility and the v2.0 eggNOG database (18).

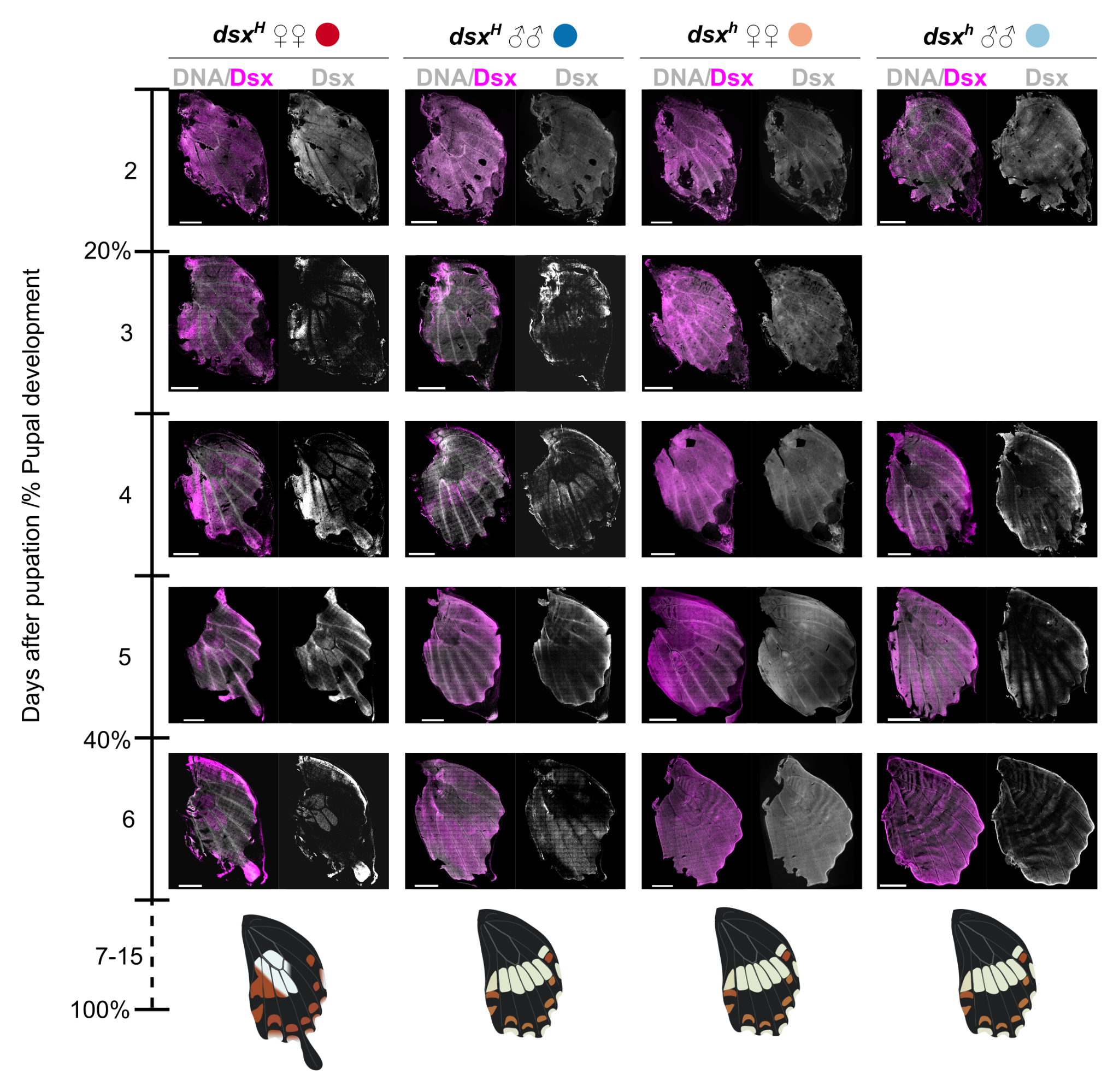

**Fig S1. Additional anti-Dsx stains in pupal hindwings.** Related to Fig 1. Diagrams of adult color patterns are shown below. Note that intensities should only be compared across any individual wing, not between wings. Saturated regions (such as the tip of the tail of the P6 mimetic female) are typically due to over-fixation caused by residual membrane before staining. Scale bars: 2 mm.

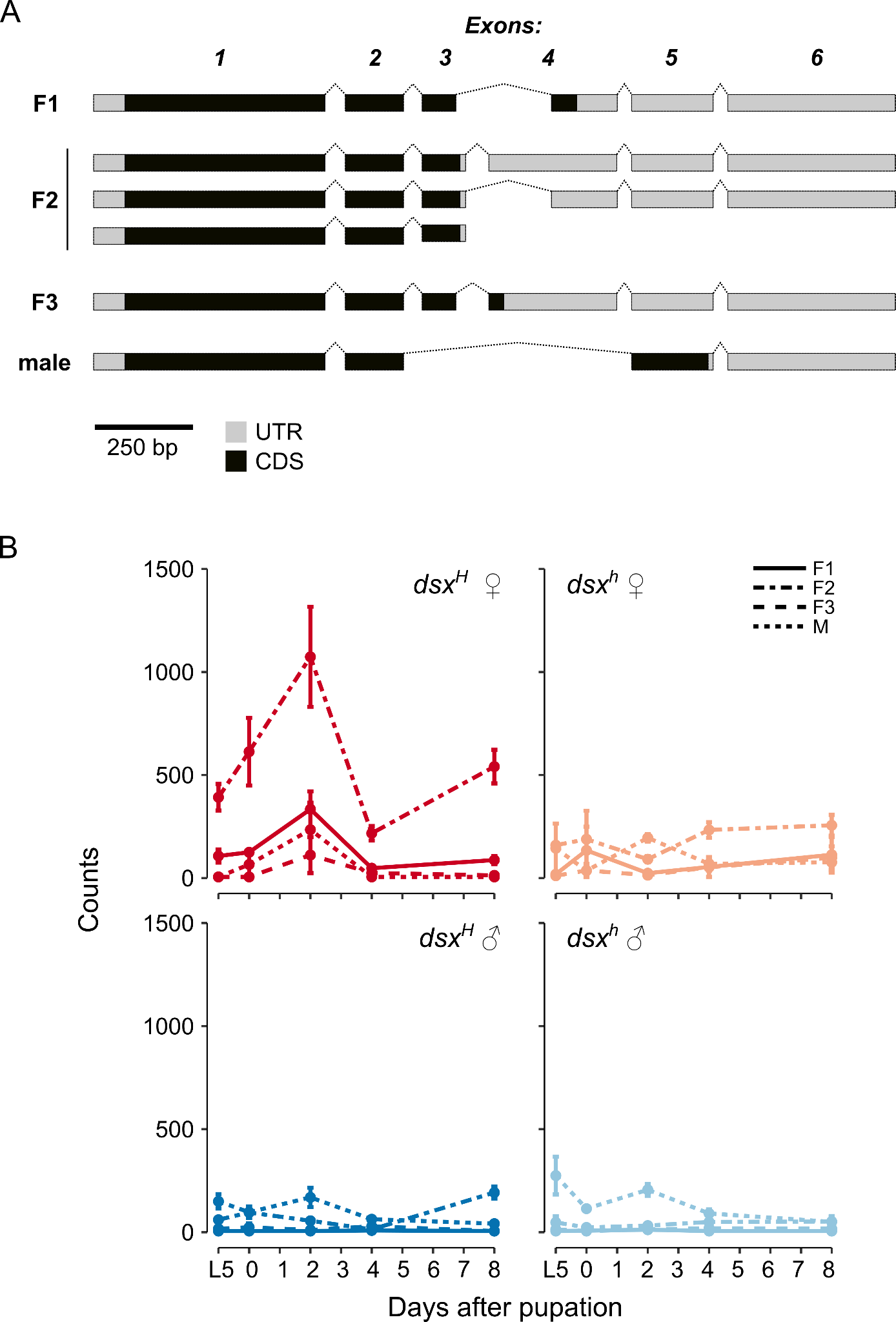

**Fig S2. *doublesex* isoforms and expression.** Related to Fig 1. (A) Exon-intron structures of the various isoforms. Introns are not to scale. (B) Normalized transcript expression levels. Previous cDNA cloning and Sanger sequencing did show that all six major isoforms are present in male and female pupal hindwings ((3) and A. Tenger-Trolander, unpublished data)

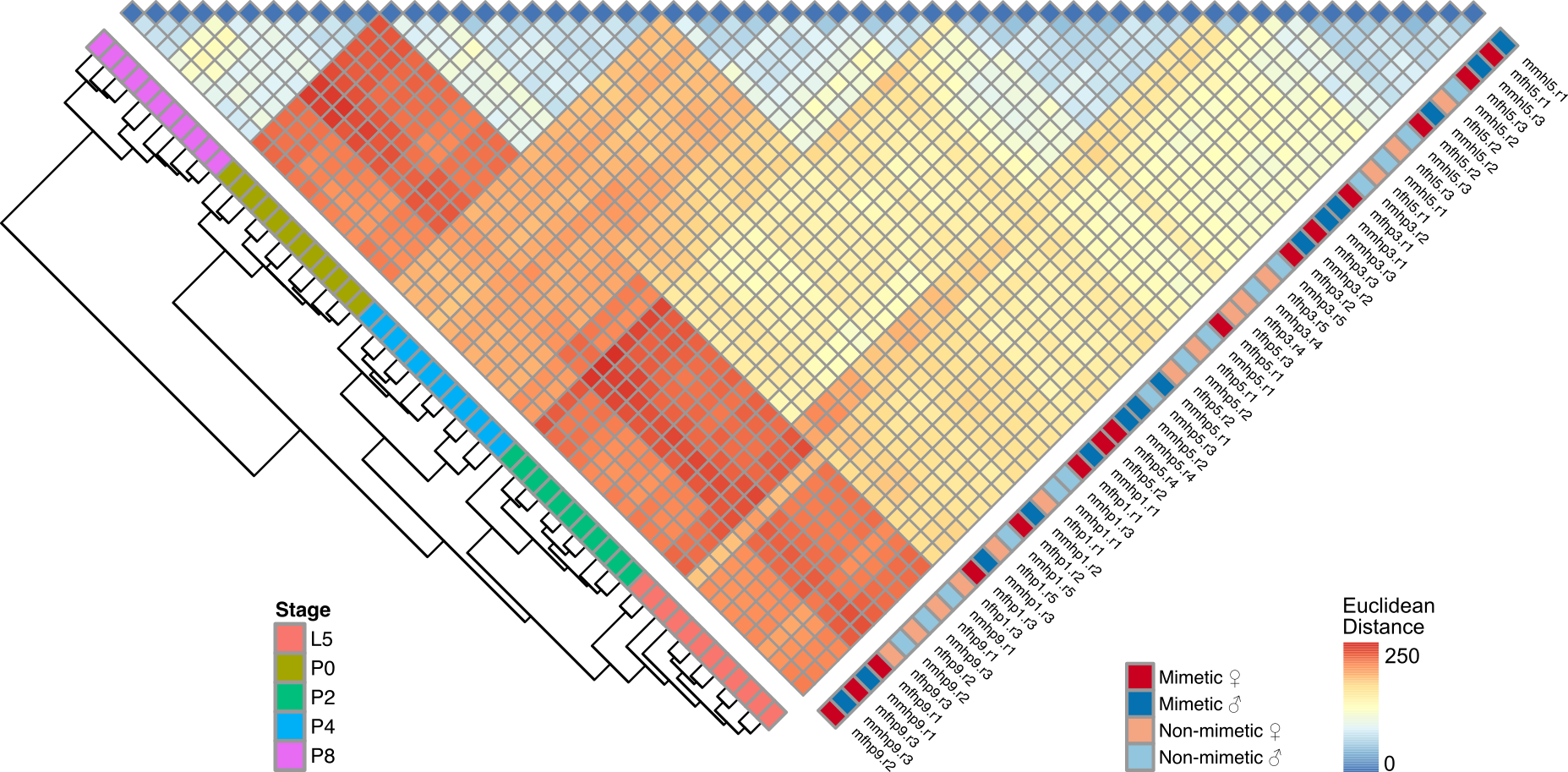

**Fig S3. Euclidean distance heatmap of all RNA-seq samples based on normalized expression levels of 12,721 genes.** Samples were clustered using the pheatmap package and *k* means clustering (19).

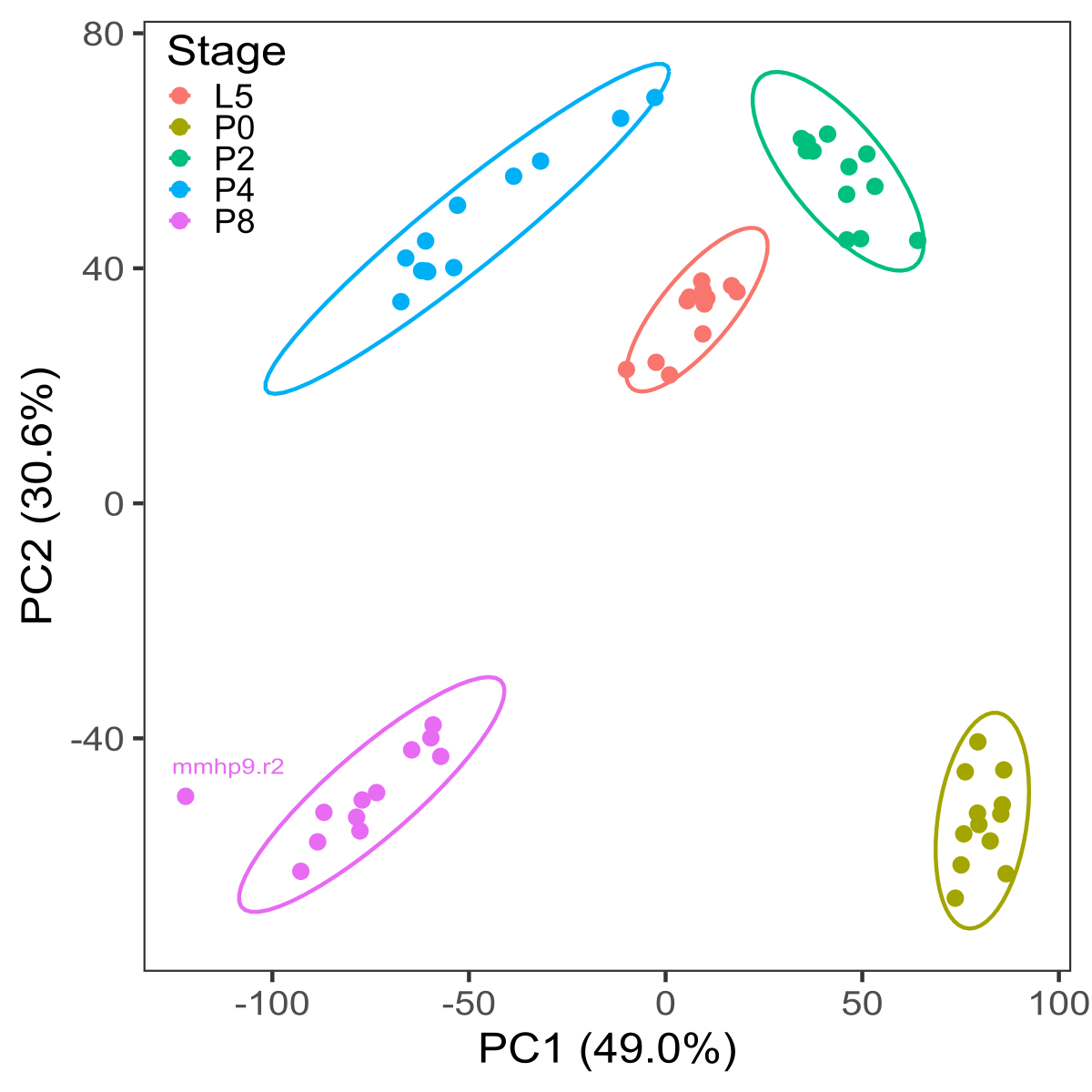

**Fig S4. Robust PCA of all 60 RNA-seq samples based on normalized expression levels of 12,721 genes.** Ellipses show 95% confidence intervals based on the multivariate *t* distribution, implemented in the rrcov R package (20).

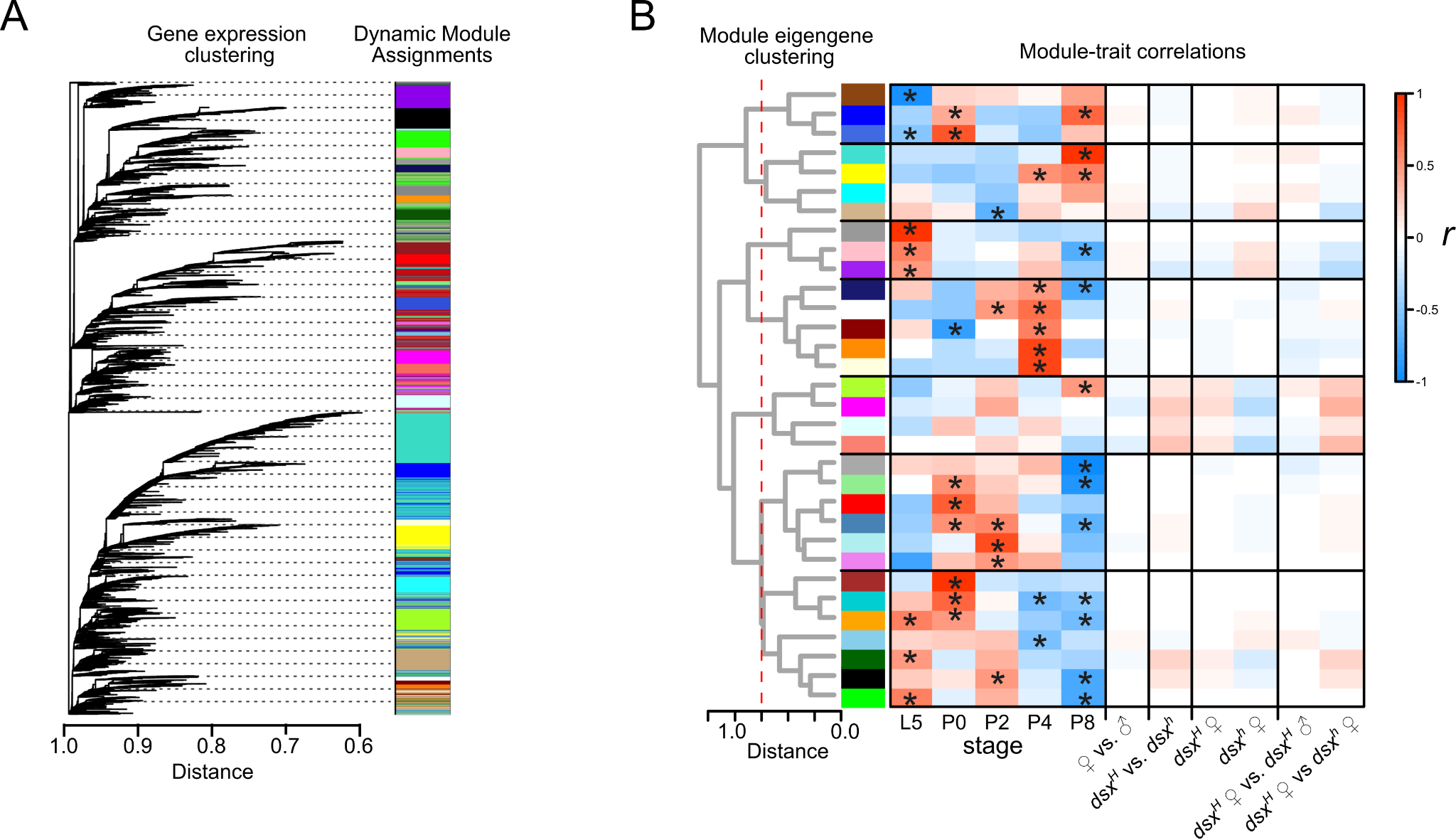

**Fig S5. Gene co-expression network construction and analysis using WGCNA.** A) Clustering and module assignment. B) Module eigengene clustering, meta-module identification, and correlations between module eigengenes and select comparisons. *: Bonferroni-corrected *p* < 0.05.

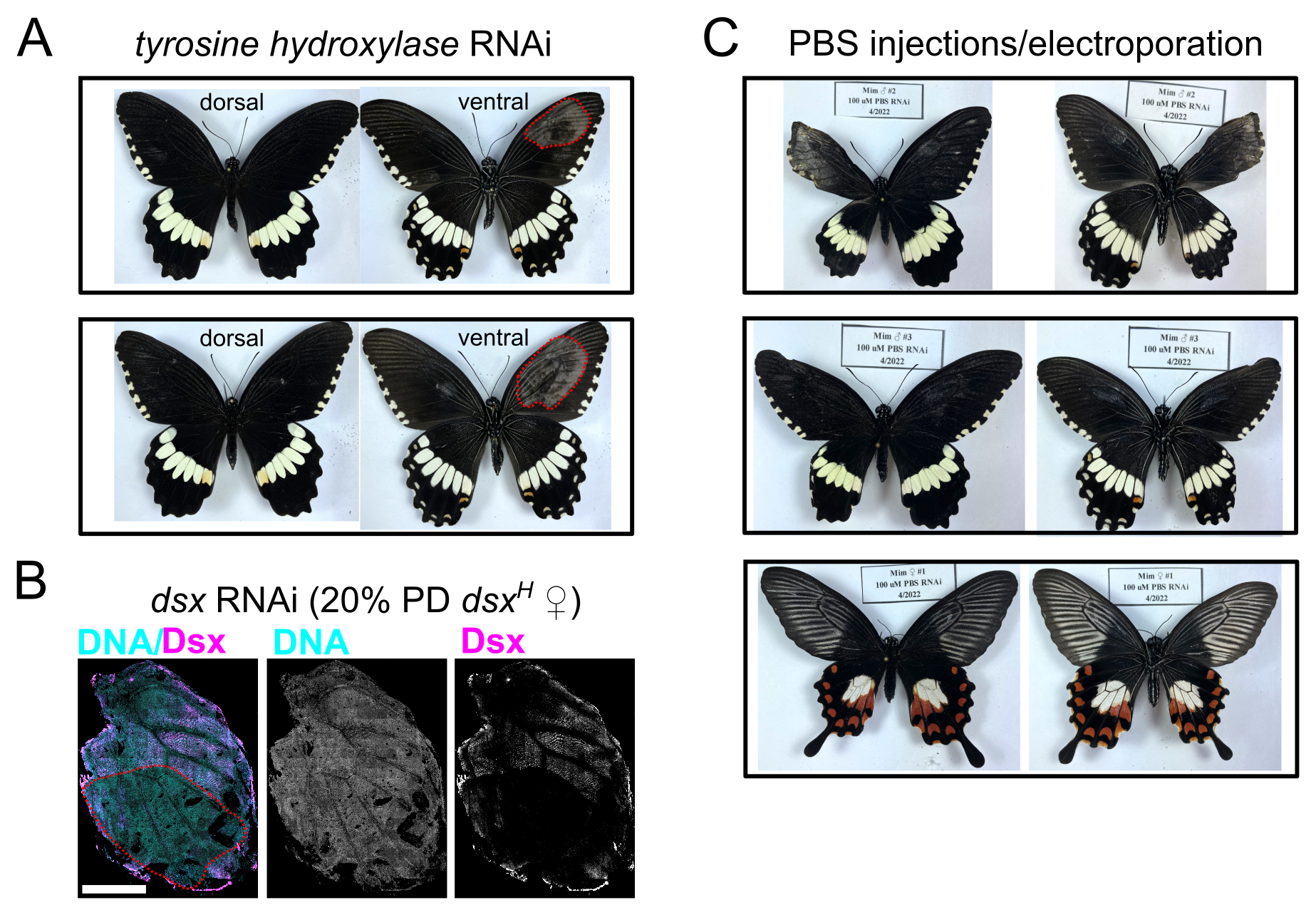

**Fig S6. RNAi injection controls.** A) Knockdowns of TH, a crucial gene in the melanin synthesis pathway that does not turn on until ~12 days after pupation (when the wing is injected) (21). DsiRNAs were injected into the left forewing and electroporated as described in the Methods. These are both *dsx^H^* males. B) Anti-Dsx staining of a mimetic female hindwing injected with DsiRNA against *dsx*. Compare to Fig 1 and Fig S1 at 20%PD, where you see the expected Dsx staining pattern across the distal 75% of the wing. C) Negative controls. We electroporated 1.5 uL PBS (DsiRNA carrier solution) into the left ventral hindwing.

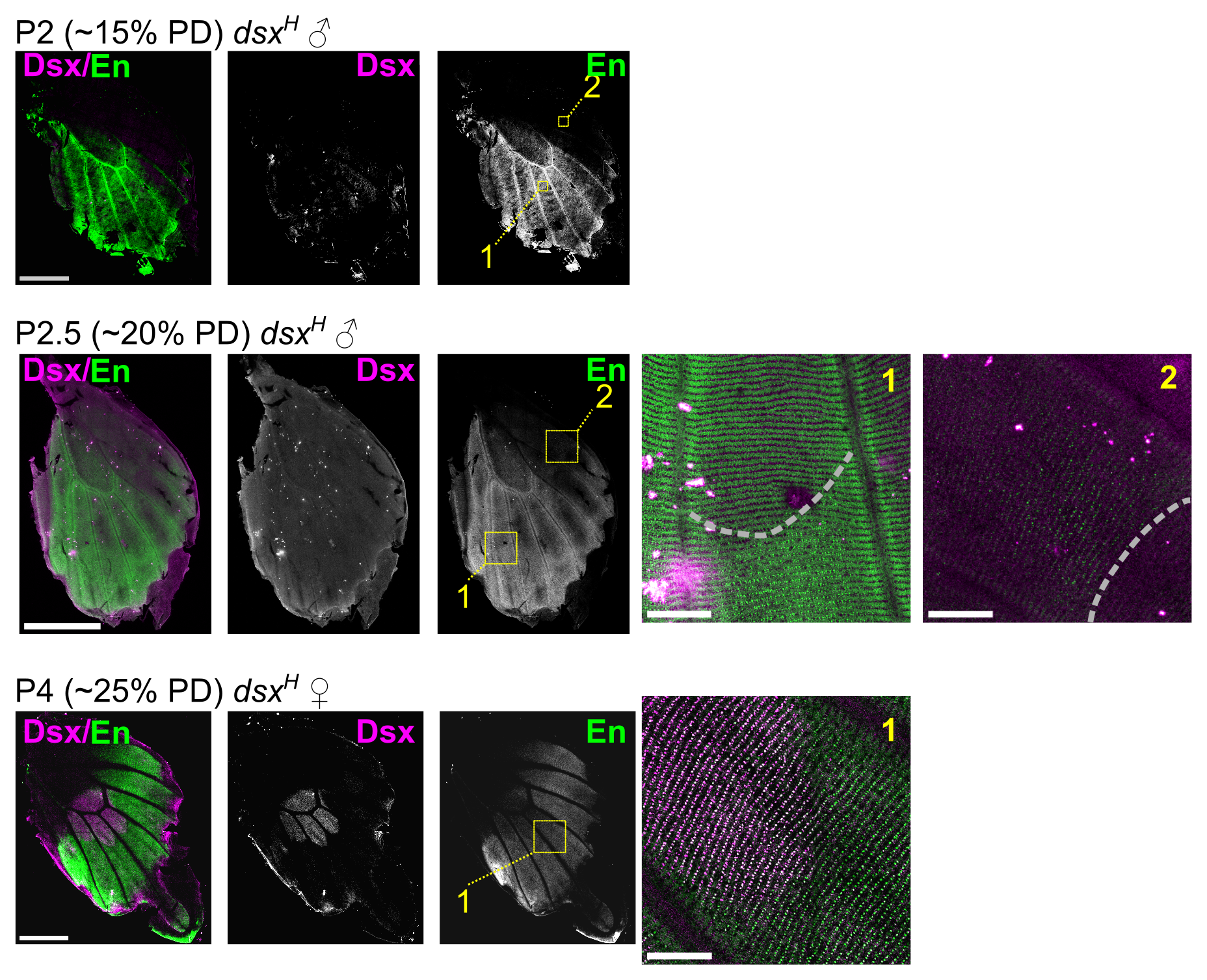
**Fig S7: Additional anti-En stains.** Zoomed images were taken with the same settings and not adjusted, so they are comparable. Scale bars: 2 mm for full wings, 200 um for zooms.

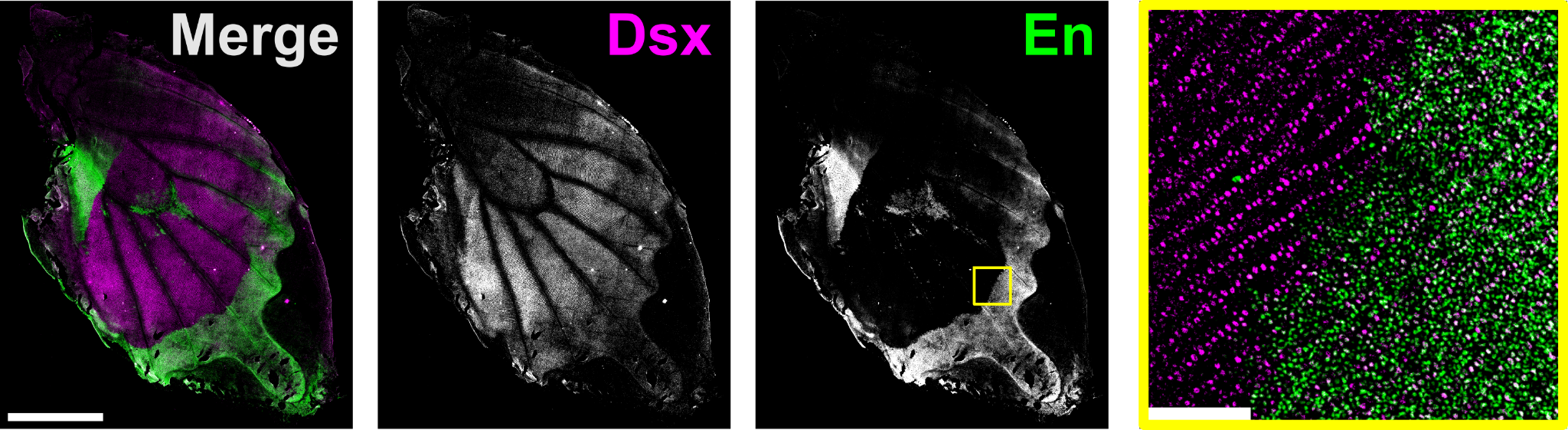

**Fig S8. *en* RNAi does not affect Dsx expression.** We injected 1.5 uL of 100 uM *en* DsiRNA into the wings between Cu1 and M3 near the discal cell and electroporated as described in the methods. Pupa was allowed to develop for three days before dissection and staining. Compare to Fig 1C mimetic female 25% PD and Fig S1 mimetic female P3 - P4.
