## Supplementary Dataset for "Conserved signaling pathways antagonize and synergize with co-opted *doublesex* to control development of novel mimetic butterfly wing patterns"

**RNAi tests of potential Dsx targets**

NWV, 7-2021 through 08-2022. Injected 1.5 uL 100 uM DsiRNAs in water into the left hindwing, shocked 5 x 0.28 s (10 V), with cathode ventral and anode dorsal (DsiRNA should move into ventral cells). We aimed to inject ~3 males and ~3 females for each genotype (homozygous non-mimetic *dsx*, homozygous mimetic *dsx*).

Below are images of each insect and summaries of the phenotypes. NND: no noticeable difference between control and injected wings.

Landmarks, including veins and the outlines of colored patches. Mimetic “white” patches are truly unpigmented white scales. Non-mimetic light patches are pale yellow and are pigmented.

- **Marginal crescents** abut the distal edge of the wing.
- **Sub-marginal crescents** (SMCs) are just inside the distal margin.
- The **A-Cu2 black spot** is bordered by red scales from the A-Cu2 red patch and **anal triangle** to varying extents and frequently make what looks like a hook.

There are essentially six types of scales:

- **Melanic**
- **Red**
- **White** (mimetic)
- **Pale yellow** (non-mimetic, males)
- **Opal** (iridescent, pale blue) – scattered throughout red regions, rarely in melanic regions
- **Light orange** – scattered throughout melanic regions

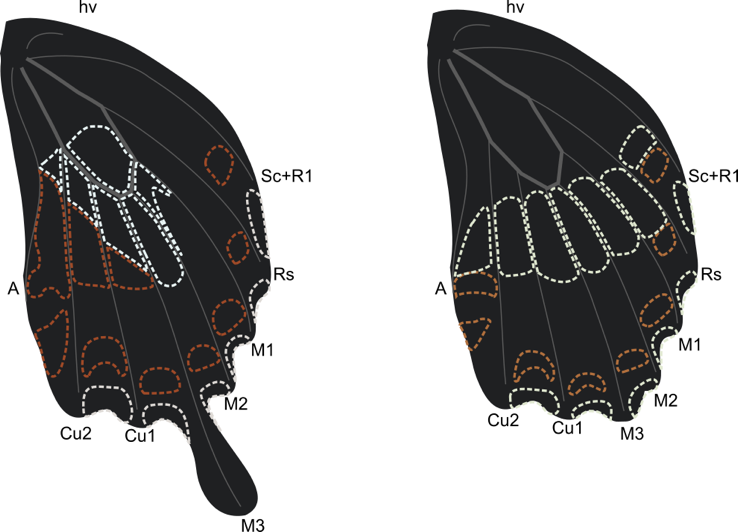

Diagram of Papilio hindwing veins and *P. polytes alphenor* color pattern elements.

***PBS (negative control)***

These injections tested the effects on color pattern of PBS, where we expect no phenotype.

**Phenotype summary**:

We saw NND.

**Mimetic Females**

**MF1:**

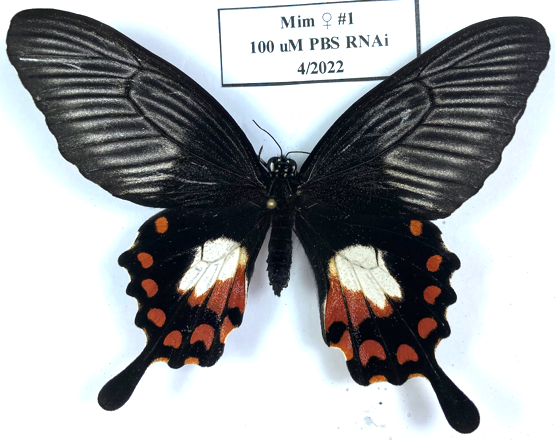

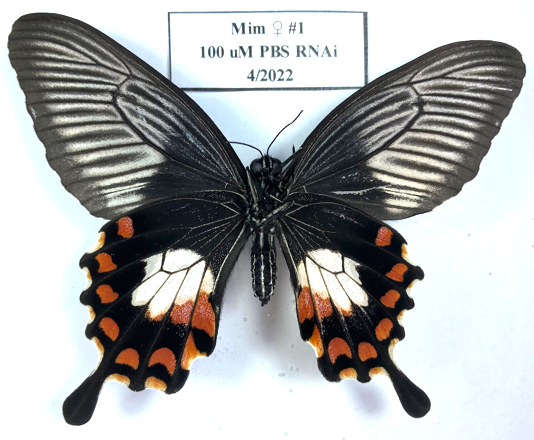

Dorsal:

- NND

Ventral:

- NND

**Mimetic Males**

**MM2:**

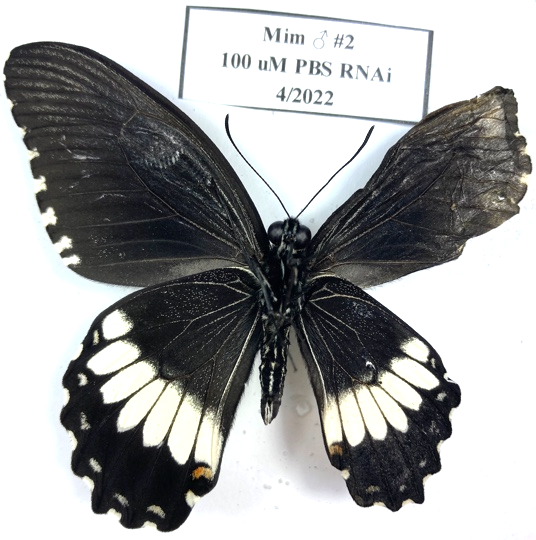

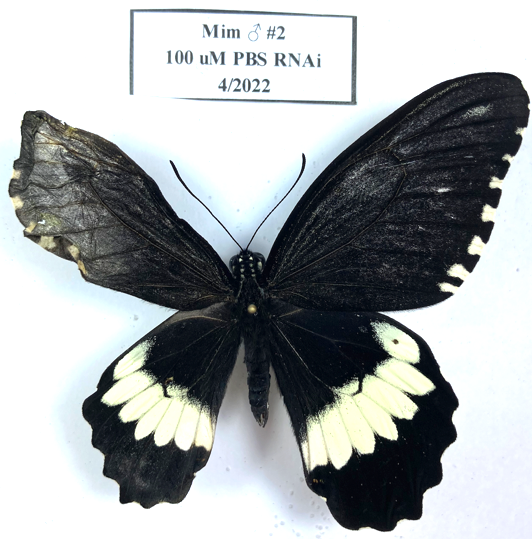

Dorsal:

- NND beyond some damage to wing shape during emergence. Wings get stuck occasionally, pulled out, then dry wrinkled.

Ventral:

- NND

**MM3:**

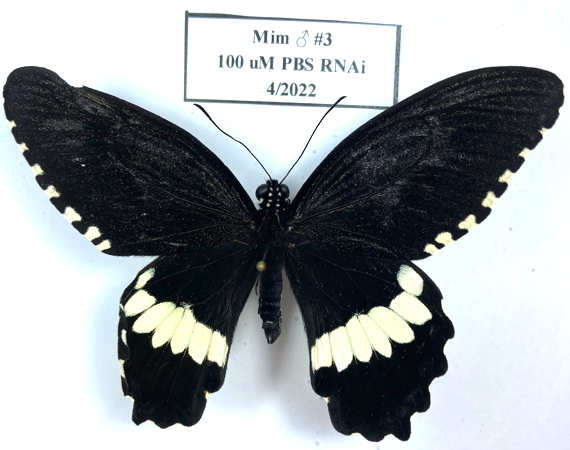

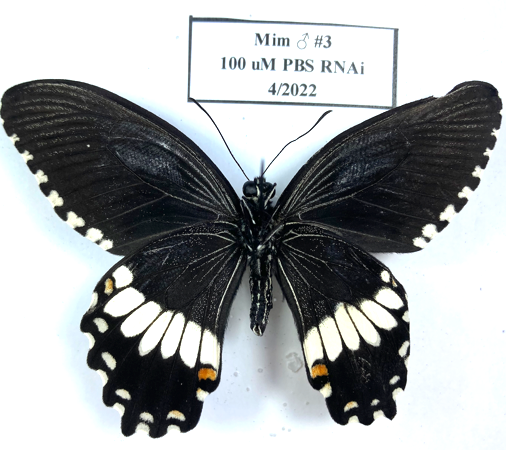

Dorsal:

- Slight damage to proximal edge of Cu1-M3 pale patch, small patch of pale scales.

Ventral:

- Slight damage to proximal edge of Cu1-M3 pale patch, small patch of pale scales.

***evm.TU.chr18.639***

The gene *evm.TU.chr18.639* encodes a protein with no known orthologs. BLASTp to the nr database yields hits only to hypothetical proteins from moths. The closest hit (e-value 0.74) in *D. melanogaster* is to Notch.

We had no expectations for phenotypes induced by *evm.TU.chr18.639* RNAi.

***evm.TU.chr18.639 expression***

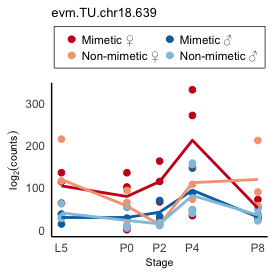

**Phenotype summary**:

Results suggest that *evm.TU.chr18.639* acts either to suppress red scale formation or promote melanic scale formation along the Cu2 vein between the red patches and AT or SMC specifically in mimetic females. Melanic appears to be the default fate. It is possible that this gene also promotes opal scale formation, but opal and red scale development are co-dependent (i.e. opal scales primarily develop in red regions, especially the A-Cu2 and Cu2-Cu1 red patches). *chr18.639* knockdowns in butterflies with non-mimetic color patterns results in a small or no increase in red scale frequencies in Cu2-Cu1 SMC or along the Cu2 vein between the A-Cu2 red patch and AT.

**Mimetic females**

**MF1:**

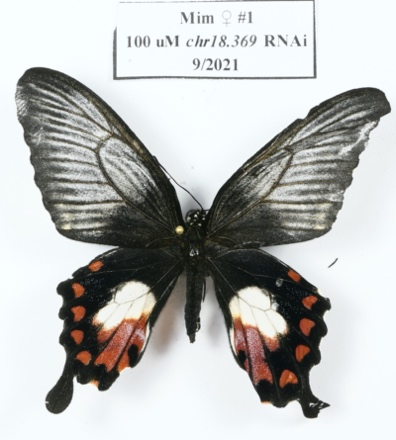

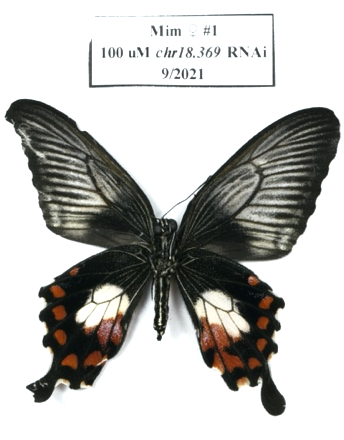

Dorsal:

- Cu1-Cu2: increased red scales, connecting red patch and SMC

Ventral:

- A-Cu2 and Cu2-Cu1: expanded red patch (along Cu2 vein) connecting patch to AT or SMC.
- A-Cu2 and Cu2-Cu1: reduced frequency of opal scales in red regions

**MF2:**

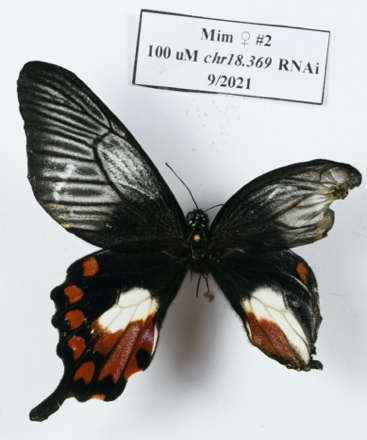

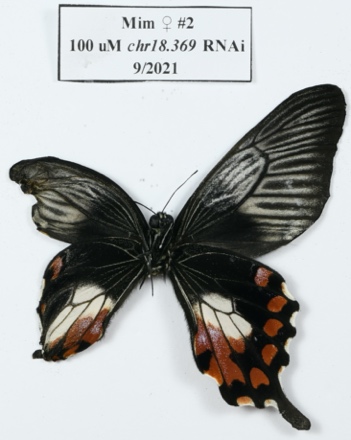

Phenotypes hard to tell because control wing was damaged during emergence.

**MF3:**

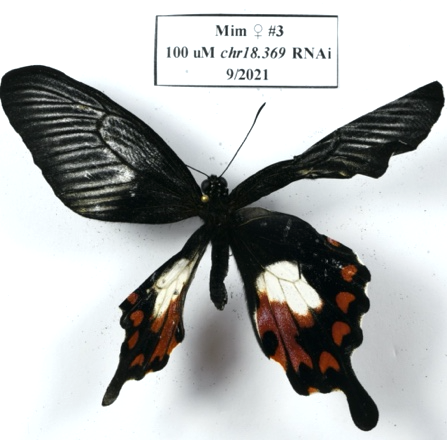

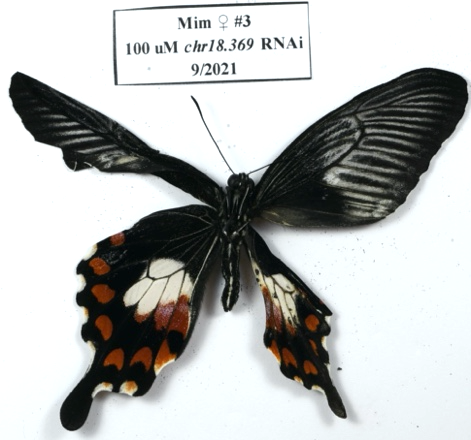

Tough to tell because injected wing is crumpled.

Dorsal:

- Cu2-Cu1: increased red scales, filling melanic patch between red patch and SMC.
- Cu2-Cu1: Reduced frequency of opal scales?

Ventral: unknown

**MF4:**

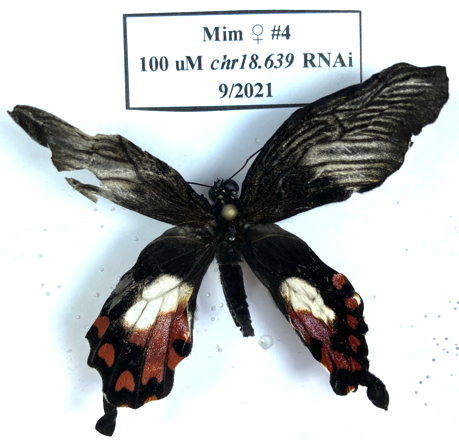

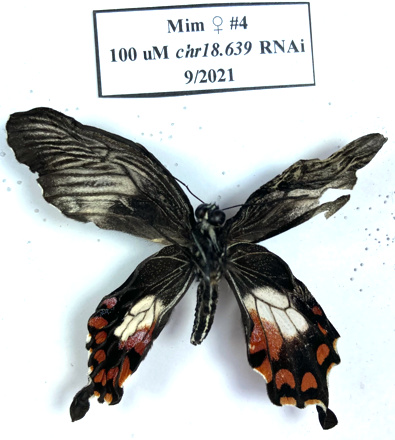

Dorsal:

- A-Cu2: Increased red scales along Cu2 vein, completing hook/black spot
- Cu2-Cu1: Increased red scales along Cu2 vein, completely merged red patch with SMC

Ventral:

- A-Cu2: Hook/black spot completed
- Cu2-Cu1: anterior expansion of red patch

**Mimetic males**

MM1:

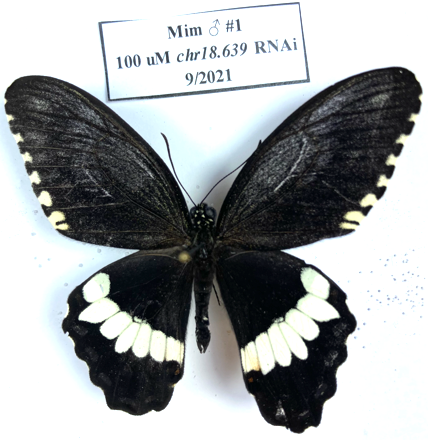

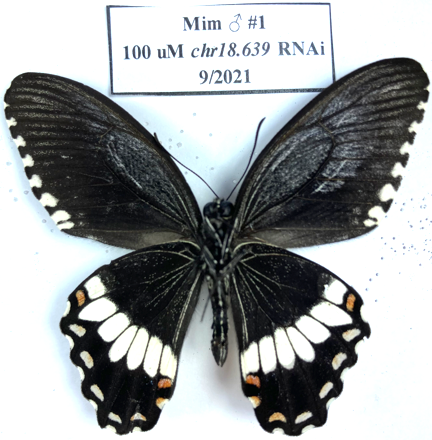

Dorsal:

- A-Cu2: significantly fewer orange scales in white patch

Ventral:

- Cu2-C1: significantly more red scales in SMC

**MM2:**

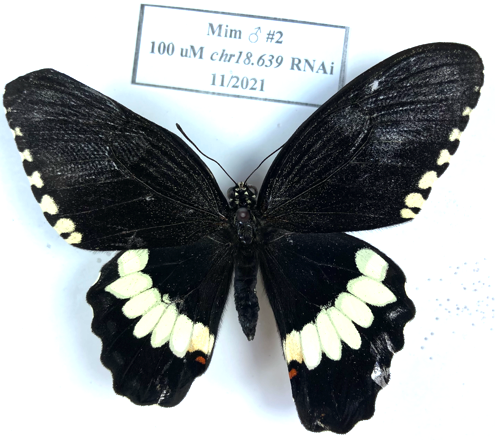

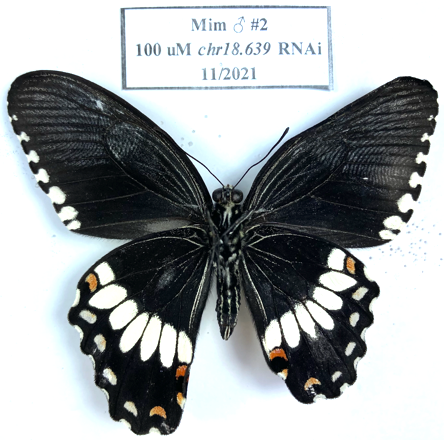

Dorsal:

- Pale scales in DC

Ventral:

- Irregular posterior edge of Cu1-M3 white patch, grayish scales distal to white patch – could be damage?

**MM3:**

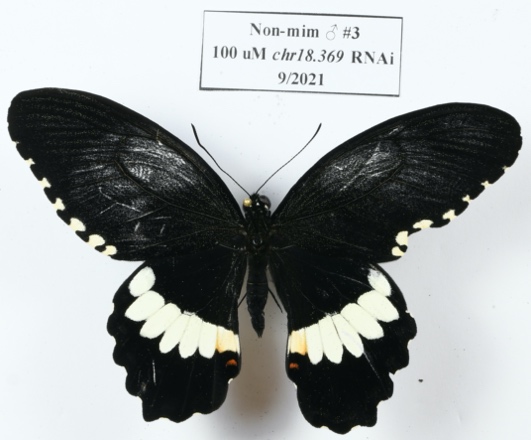

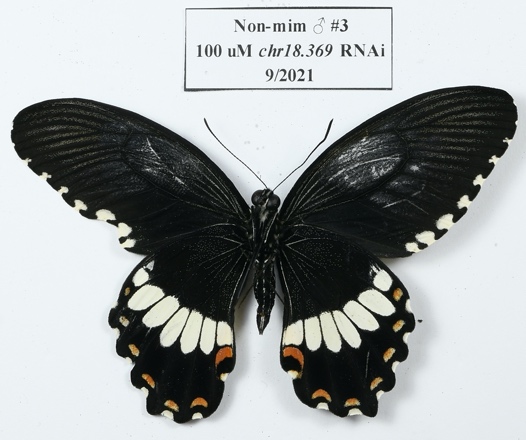

Mislabeled before taking pictures. Fixed now.

Dorsal: NND

Ventral:

- A-Cu2: Slightly more red scales between red patch and AT.

**Non-mimetic Females**

**NF1:**

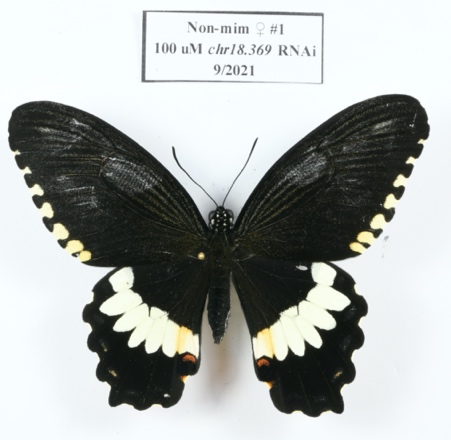

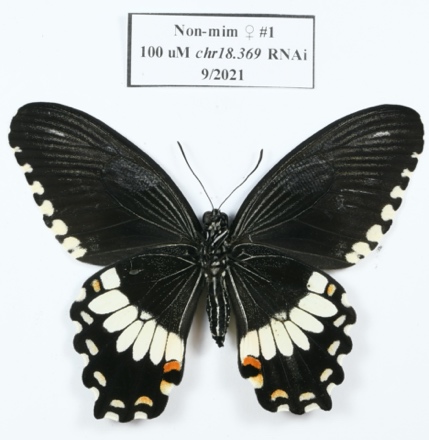

Dorsal:

- NND

Ventral:

- Slight increase in melanic scales in discal cell white patch

**Non-mimetic Males**

**NM1:**

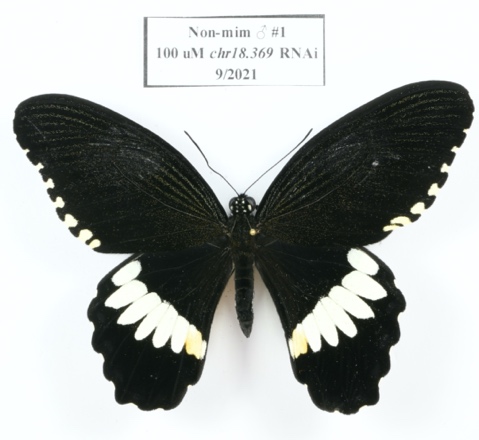

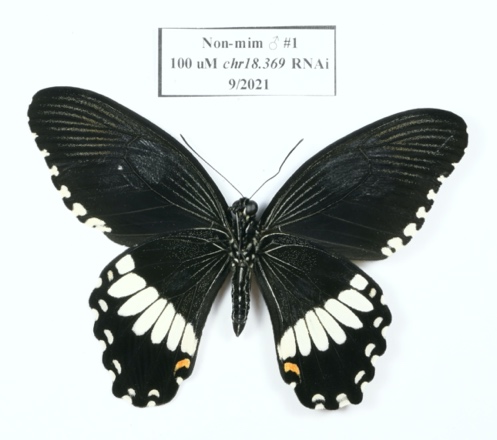

Dorsal:

- NND

Ventral:

- NND

**NM2:**

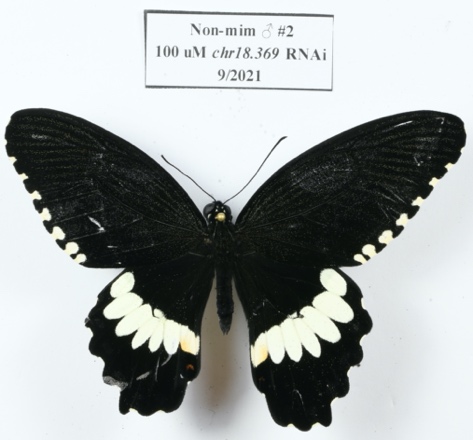

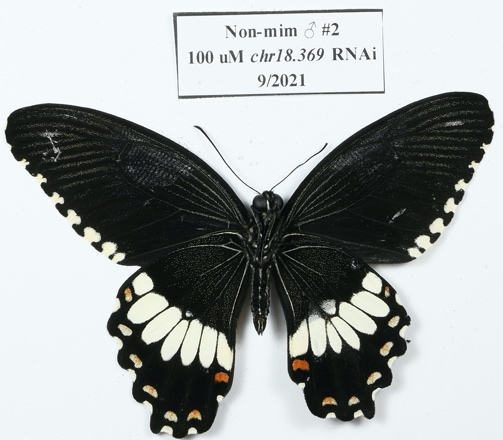

Dorsal:

- Slightly larger A-Cu2 SMC

Ventral:

- Cu2-Cu1, Cu1-M3, DC: anterior expansion of pale patch, appears to be gray scales. **Could be damage from the injection.

***WntA***

*WntA* (*evm.TU.chr4.196*) is known to specify boundaries between melanic and colored patches across butterflies (Mazo-Vargas et al., 2017, PNAS), albeit in a variety of ways. This gene is most similar to fly *Wnt2* and frequently labeled as Wnt4-like (less frequently Wnt1-like) in other non-model organism genomes. *WntA* is a soluble ligand for the *frizzled* receptor and activates canonical and conserved signaling pathways leading to changes in gene expression controlled by *β-catenin* (fly *armadillo*).

I expected to see changes to the melanic/non-melanic boundaries, but did not know where those changes may be.

***WntA expression***

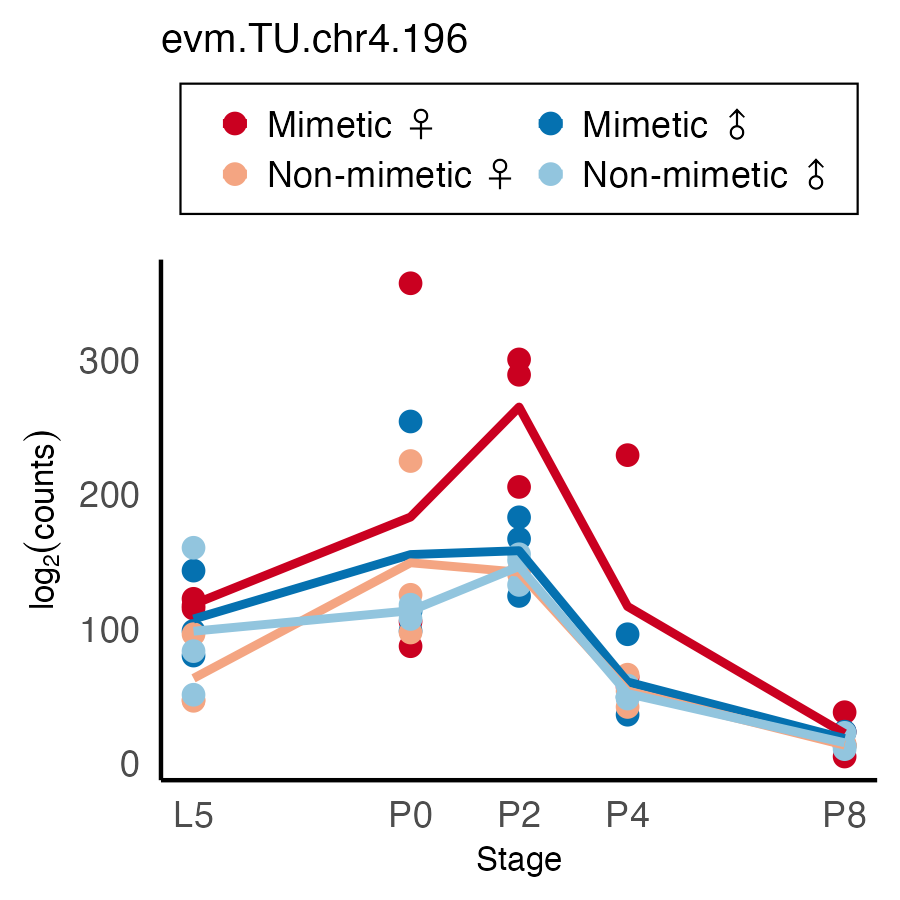

**Phenotype summary:**

Consistent with other classes of phenotypes observed in nymphalids, *WntA* generally establishes the posterior boundaries of white/pale yellow regions. Knockdowns in both mimetic and non-mimetic females resulted in extensive posterior expansion of these white/pale regions near the injection site, but no anterior extension. It is not completely clear to me if the white patches are only expanding to the edge of the electroporation bubble, but I think that this is not the case for a couple reasons. First, the white never expands into the SMCs – the anterior boundaries on the SMCs are retained even in the strongest knockdowns. Second, the A-Cu2 red patch and AT (and the melanic spot they surround) are preserved in mimetic females. Furthermore, mosaic knockdowns (e.g. MF4) show that the red pattern is unchanged and that white scale fate overrides the red, opal, and melanic scale fate when *WntA* is knocked down.

These knockdowns establish the scale fate hierarchy as: white > red / opal > black. *Although see *en* RNAi – regions that are normally red are comprised almost solely of opal scales. The existence of some immutable color patterns is different from what Mazo-Vargas et al. (2017) showed, and actually suggests that *WntA* may be specifying the distal boundary of the pale/white patches rather than the positions of the melanic regions.

These knockdowns further establish that the SMCs and the A-Cu2 red patch and AT (and the melanic spot they define) are specified by a separate mechanism. The A-Cu2 pattern appears to be larger and more strongly defined in mimetic females than non-mimetic females. *See also *en* RNAi.

**Mimetic Females**

**MF1:**

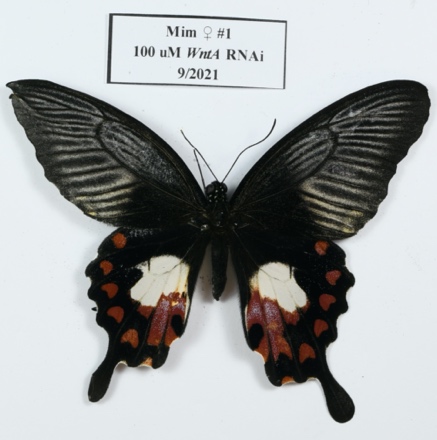

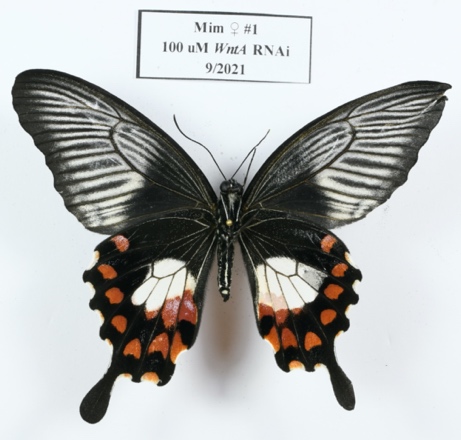

Dorsal:

- A-Cu2: posterior expansion of white region, some mosaicism in red patch. Flat posterior edge on SMC (which remains red).
- Cu2-Cu1: ~10% posterior expansion of white patch, increased white scales in marginal spot
- Cu1-M3: ~10% posterior expansion of white patch
- M3-M2: ~10% posterior expansion of white patch

Ventral:

- Posterior expansion of white patches in medial 5 cells. Red patterns appear unaltered. White expansion overrides red patches. Most noticeable in M2-M1 cell.
- **The white is still mimetic white. You can tell the difference between white and pale yellow scales visually, and especially when you shine UV light on them (pale fluoresce).
- **Unclear if the posterior edge stops here because this is the extent of WntA action, or if it is the edge of the bubble used during electroporation. Probably the latter – see additional bugs.

**MF5:**

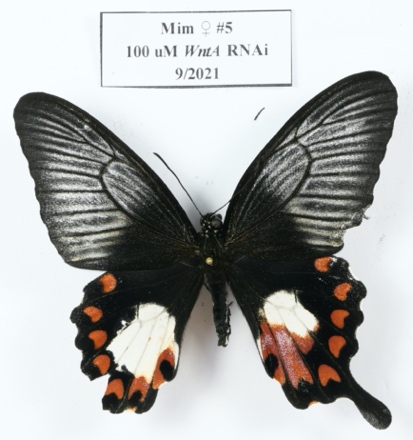

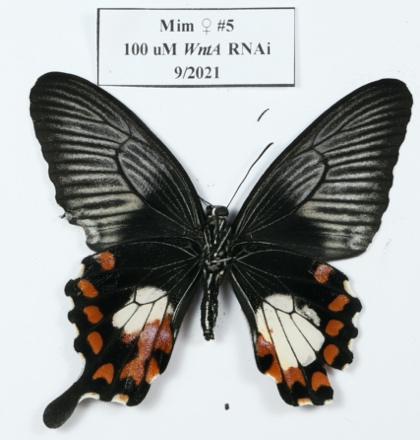

Dorsal:

- Massive posterior expansion of white patches in medial 4 cells.
  - EXCEPT the hook in A-Cu2. This appears to be its own pattern element.
  - Same with the Cu2-Cu1 SMC.
- Increased white scales in Cu2-Cu1 marginal crescent.

Ventral:

- Massive posterior expansion of white patches in medial 4 cells. Mosaicism in M2-M1.
  - EXCEPT the hook in A-Cu2 and Cu2-Cu1 SMC, as with the dorsal surface.
- White scale fate overrides both opal and red scale fate.
- While it remains uncertain whether the posterior edge of the mutant patch is defined by the bubble or *WntA* action, it seems like it’s the latter because the anterior margin of the SMC in Cu2-Cu1 is essentially normal, same with A-Cu2 red patch.

**MF4:**

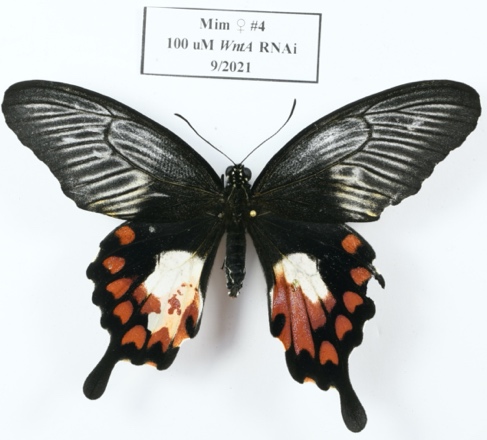

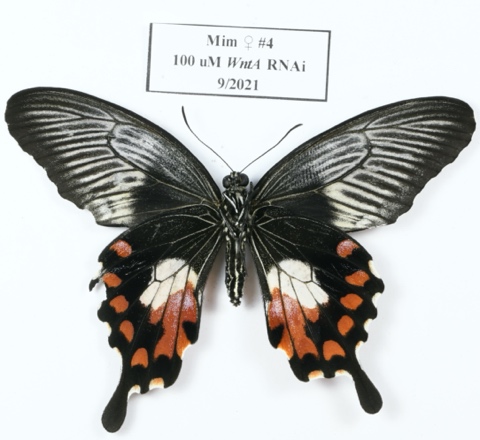

Dorsal:

- Mosaic posterior extension of white in medial 3 cells
  - EXCEPT anterior edge of Cu2-Cu1 SMC
  - EXCEPT A-Cu2 patch + AT region

Ventral:

- Slight posterior expansion of white patches in medial 3 cells

**Mimetic Males**

NONE YET

**Non-mimetic Females**

**NF1:**

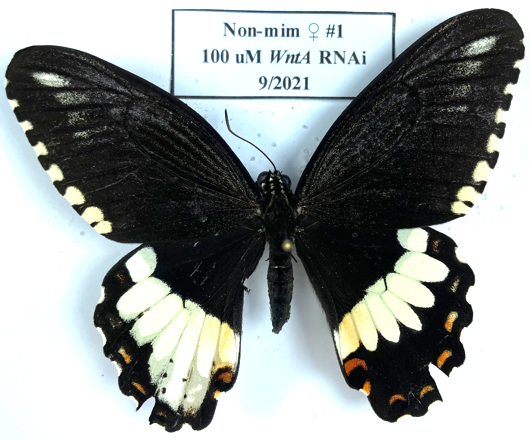

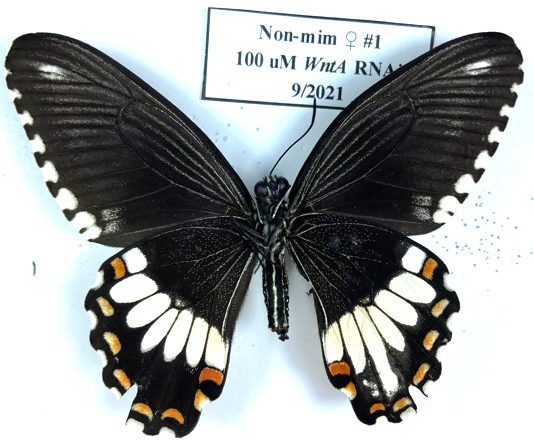

Dorsal:

- Massive posterior expansion of white patches in all cells EXCEPT Sc+R1-Rs, to the anterior edge of SMCs.
- SMCs reduced or even merged with expanded white patches in these cells
- Boundaries established by veins appear to be eliminated – smooth merges between patches in adjacent cells.

Ventral:

- Posterior expansion of white patches in Cu2-Cu1, Cu1-M3, and less so in M3-M2

**NF2:**

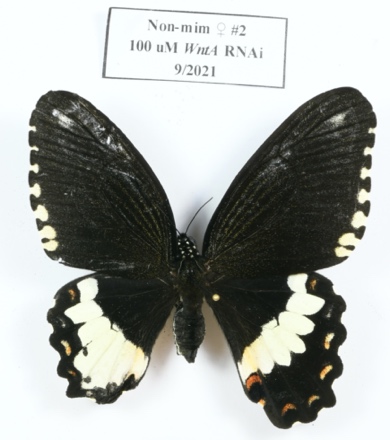

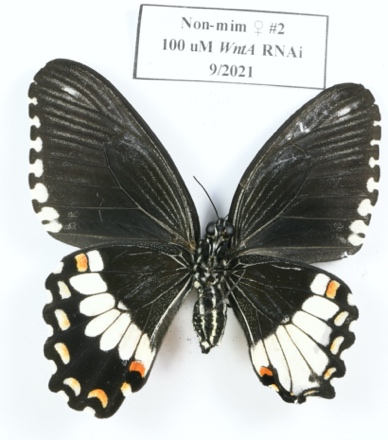

Dorsal:

- Massive posterior expansion of white patches in medial 4 cells, squishing the SMCs (but not merging with them)
- In contrast to mimetic females, the A-Cu2 red patch and AT were overridden, but the melanic spot remains.
- Boundaries established by veins appear to be eliminated – smooth merges between patches in adjacent cells.

Ventral:

- Massive posterior expansion of white patches in medial 5 cells. Slight damage to posterior edge of wing obscures exactly what’s going on there in A through M3 cells, but the SMCs do not appear to be overridden.
- More white scales in SMCs in medial 5 cells.

**NF3:**

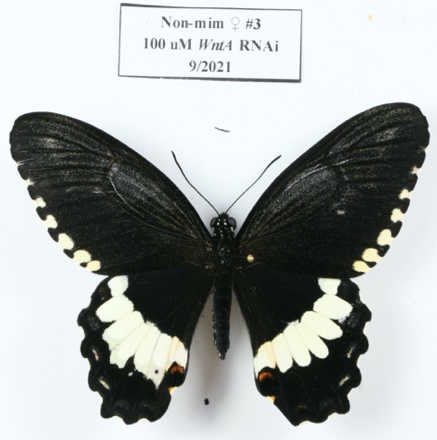

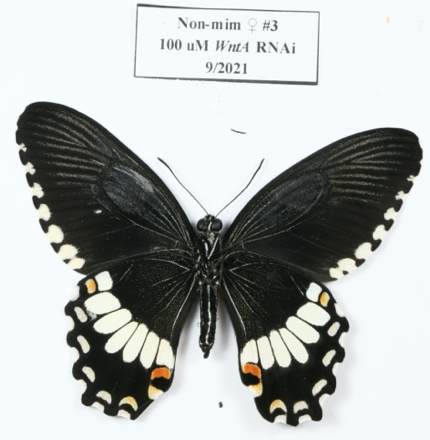

Dorsal:

- Posterior expansion of white patches in medial 5 cells
- Not as extreme as remainder of 100 uM injections, but still posterior expansion.
- Boundaries established by veins appear to be eliminated – smooth merges between patches in adjacent cells.

Ventral:

- Slight increase in white/orange scales connecting A-Cu2 red patch and AT.

**NF4:**

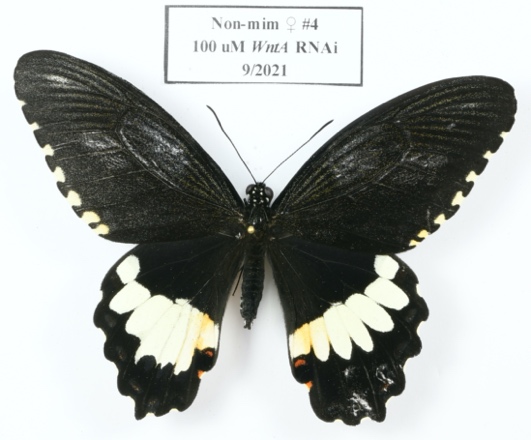

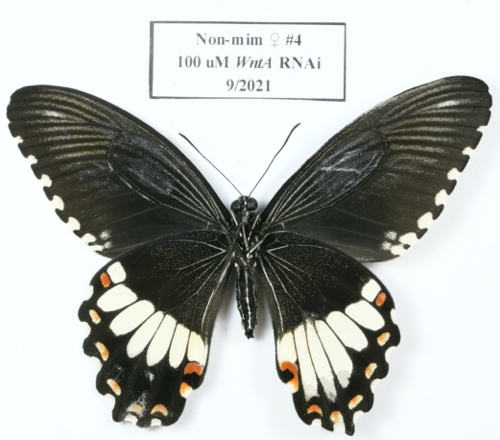

Dorsal:

- Posterior expansion of white patches in medial 5 cells. A-Cu2 red patch almost overridden.
- Boundaries established by veins appear to be eliminated – smooth merges between patches in adjacent cells.

Ventral:

- A-Cu2: Red patch and AT connected by strong stripe of red scales now.
- Posterior expansion of white patch in medial 4 cells, forcing SMCs closer to marginal crescents.
- Cu2-Cu1 expansion appears to respect anterior boundary of SMC.

**Non-mimetic Males**

**NM2:**

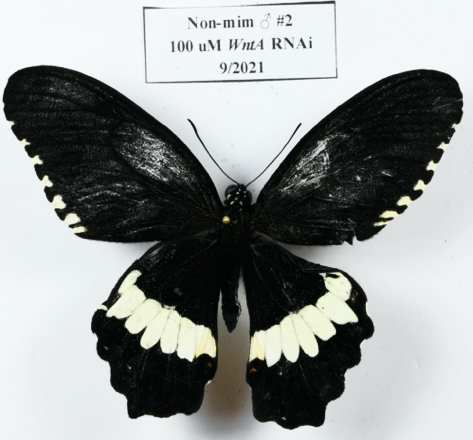

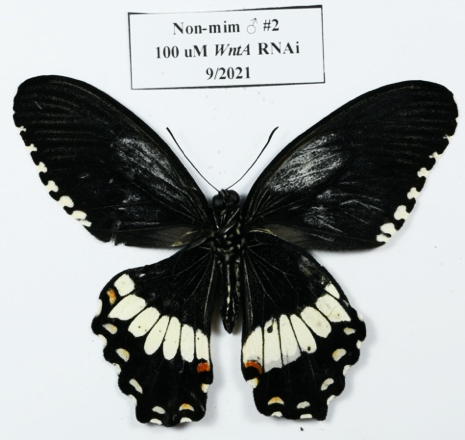

Dorsal:

- Slight extension of white patch in Cu2-Cu1.

Ventral:

- Posterior extension of white patch in medial 3 cells.
- Slight lateral expansion of red patch in A-Cu2

***GSK-3β***

*GSK-3β* (*evm.TU.chr16.419*) is the *P. polytes* homolog of *glycogen synthase kinase-3β (GSK-3β; shaggy* in *D. melanogaster*). GSK-3β is a core component of the canonical Wnt signaling pathway. This protein binds *β-catenin* in the cytoplasm, preventing it from entering the nucleus and stimulating transcription of downstream Wnt target genes. GSK-3β phosphorylates and causes degradation of cytoplasmic β-catenin. Active Wnt signaling causes *dishevelled*-mediated phosphorylation of GSK-3β, resulting in GSK-3β inactivation and release of *β-catenin*. However, GSK-3B also has a role regulating *cubitus interruptus*, so the effects could be mixed.

We expected that *GSK-3β* RNAi should increase Wnt signaling, and therefore oppose phenotypes from *WntA/Wnt1/Wnt6* knockdowns.

***GSK-3β expression***

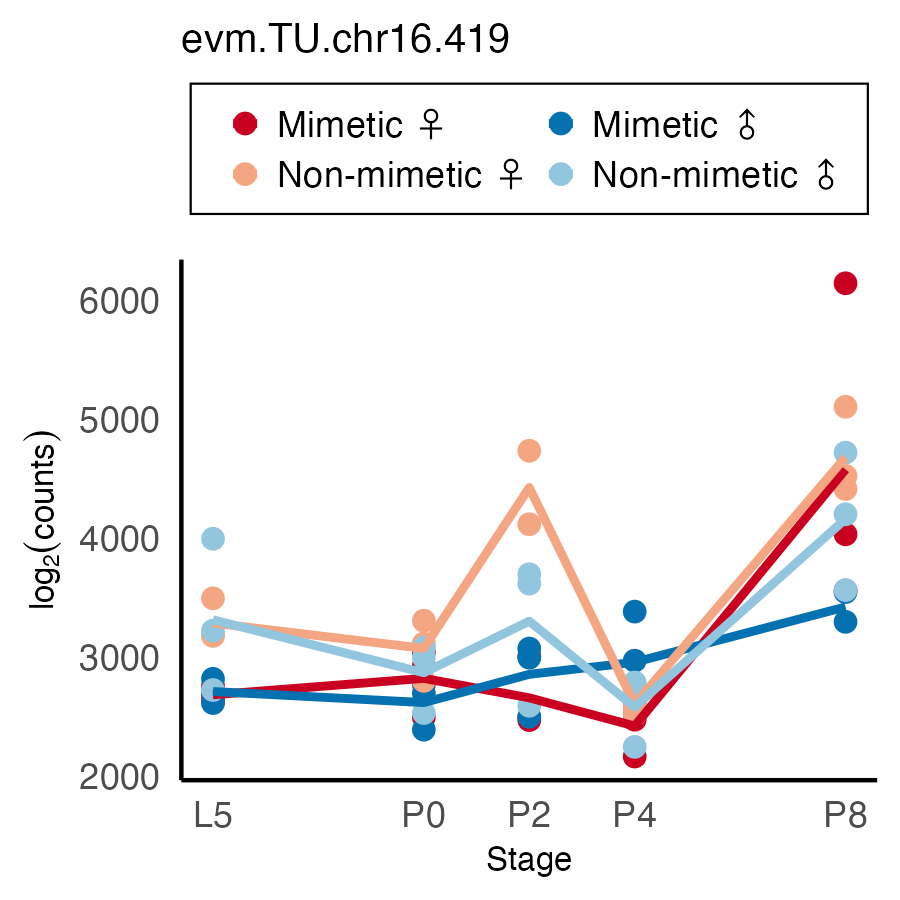

**Phenotype Summary**

*GSK-3β* knockdowns almost exclusively affect the mimetic female color patterns, particularly the distribution and frequencies of red and opal scales in the Cu2-Cu1 wing cell. All butterflies showed some blurring of boundaries between the white and adjacent red or melanic regions (i.e. some red/melanic scales pop up in the white regions near the boundary). There were also varying, but subtle effects on SMCs in bugs with the non-mimetic color pattern. These latter effects could be a general effect of Wnt signaling, because *GSK-3β* sits right in the middle of the canonical Wnt signaling pathway and Fujiwara and his colleagues have shown that Wnt1 (wg)/Wnt6 knockdowns affect those SMCs and adjacent red patterns.

**Mimetic Females**

**MF1:**

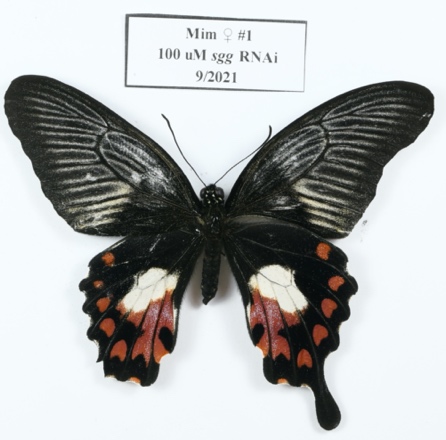

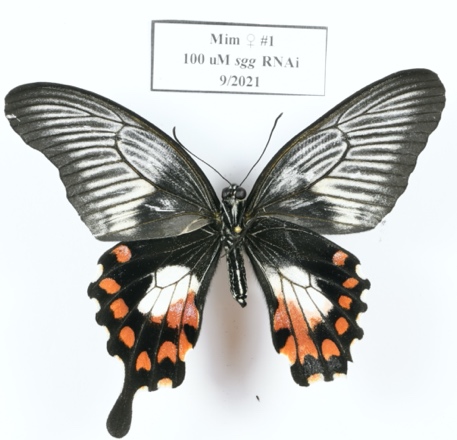

Dorsal:

- Cu2-Cu1: increased red scales connecting red patch and AT. Slight increase in red scales trailing from medial posterior point on SMC.
- M3-M2: larger red patch, seems to be because of shift of anterior edge into white patch.

Ventral:

- Reduced frequencies of opal scales in medial 4 cells, almost complete loss in Cu2-Cu1.
- Boundaries between white and red patches fuzzy in medial 3 cells.
- Slight anterior expansion of white patches in A-Cu2 and DC.
- A-Cu2: AT larger, fewer opal scales
- Cu2-Cu1: melanic region between red patch and AT filled in with red scales. Loss of opal scales.
- Cu1-M3: flat posterior boundary, red and white boundary fuzzy

**MF2:**

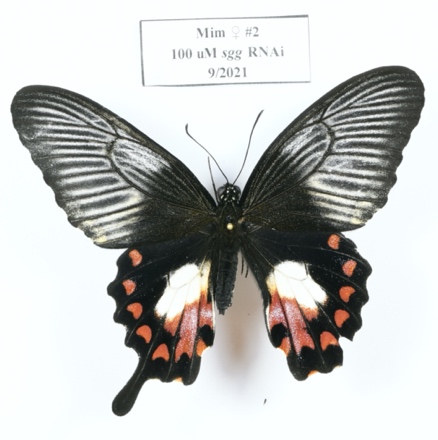

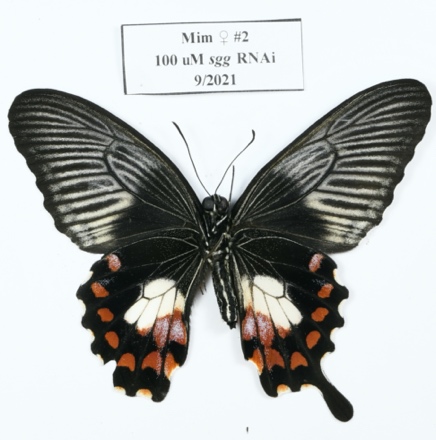

Dorsal:

- Anterior expansion of white in A-Cu2.
- Slight posterior extension of medial edge of Cu1-M3 red patch.
- Loss of opal scales in AT

Ventral:

- Anterior expansion of white in A-Cu2
- Posterior expansion of Cu2-Cu1 red patch, merging with SMC
- Reduced opal scales in Cu2-Cu1 SMC
- Very slight posterior expansion of Cu1-M3 red patch

**MF3:**

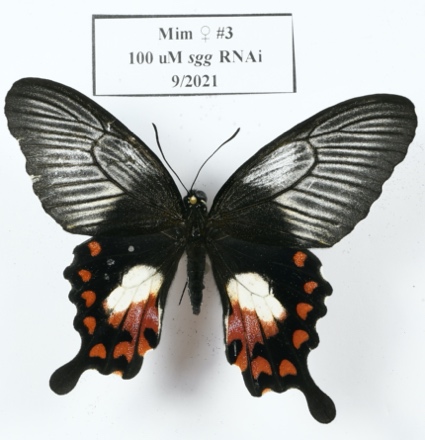

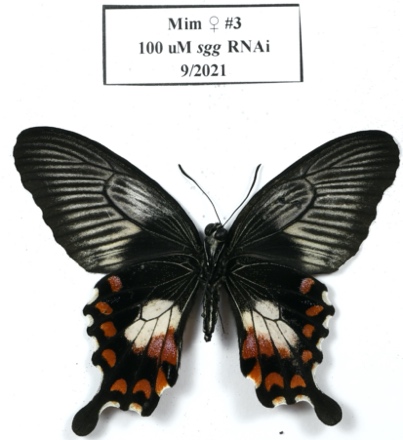

Dorsal:

- Strengthened connection (red scales) between Cu2-Cu1 red patch and SMC

Ventral:

- Posterior expansion of Cu1-M3, M3-M2 white patches into red regions, fuzzy posterior boundaries on white patches in these cells (mix of white and red/melanic scales).
- Significantly fewer opal scales in Cu2-Cu1 red patch.

**Mimetic Males**

**MM1:**

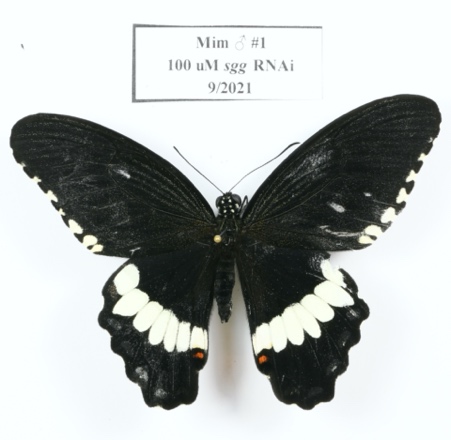

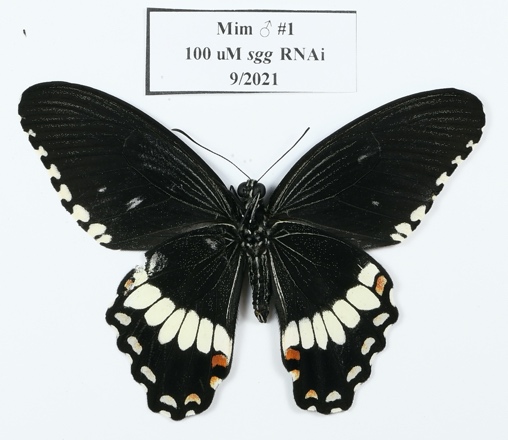

Dorsal:

- NND

Ventral:

- NND

**MM2:**

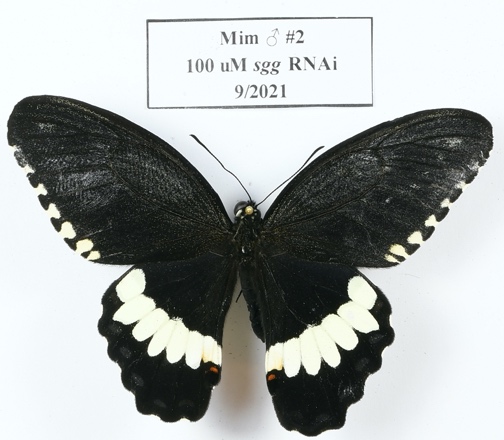

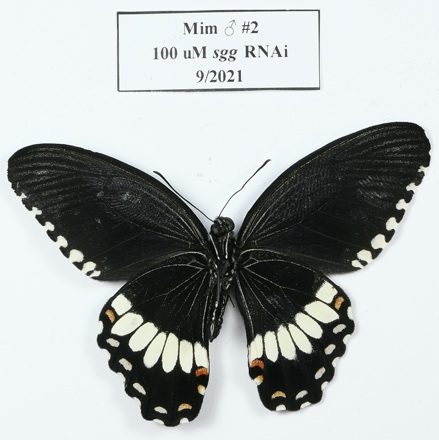

Dorsal:

- NND

Ventral:

- Slightly fuzzier posterior boundaries on Cu2-Cu1, Cu1-M3 white patches (few melanic scales and pale scales intermingled)

**Non-mimetic Females**

**NF1:**

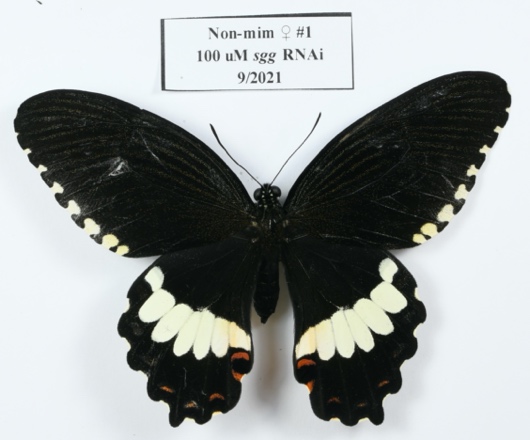

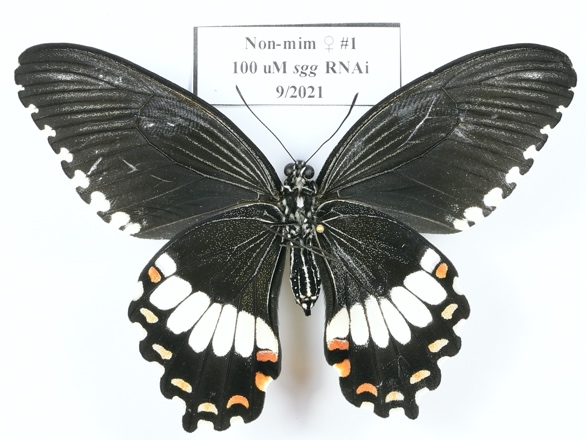

Dorsal:

- Slightly larger AT
- Slightly larger SMCs in Cu2-M2 cells

Ventral:

- NND. The increased brightness in the proximal melanic portions of the medial cells are just scales that rubbed off.

**NF2:**

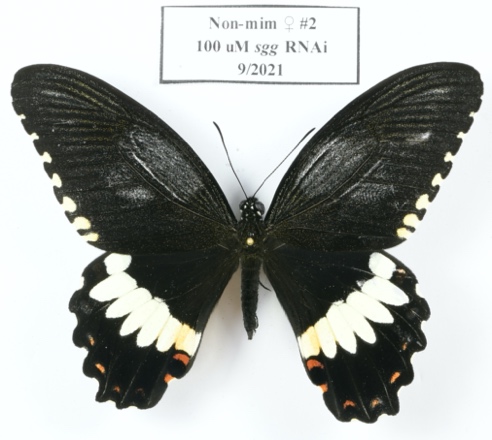

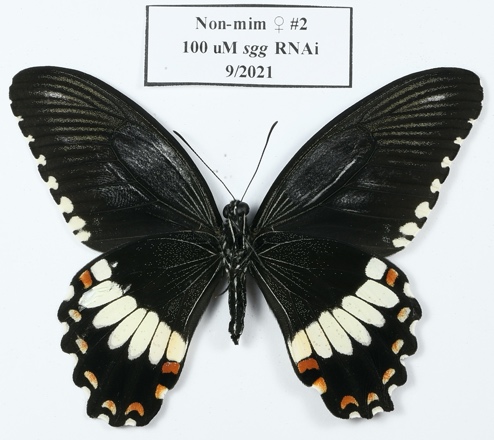

Dorsal:

- Slightly smaller SMCs in Cu2-Cu1, Cu1-M3 cells

Ventral:

- NND

**Non-mimetic Males**

**NM1:**

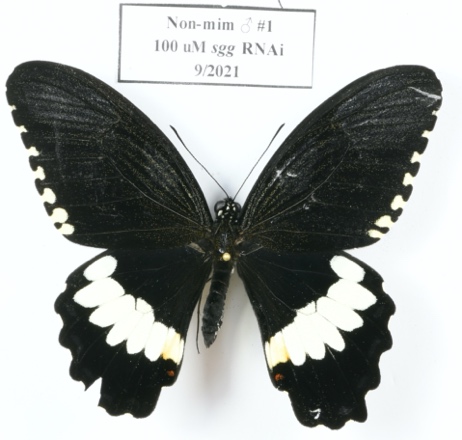

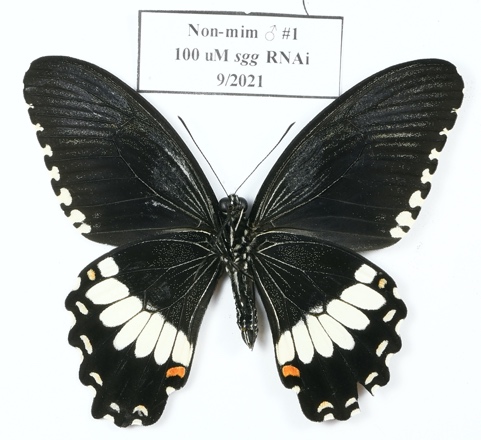

Dorsal:

- NND

Ventral:

- Slightly fuzzy anterior edge of Cu2-Cu1 and DC white patches.

***ebi***

*ebi* is a core regulator of Notch, EGFR, and Wnt signaling, complexing with histone modification enzymes and MEDIATORs to regulate chromatin (particularly histone deacetylation, as it is complexed with HDAC3), usually resulting in increased transcription of target genes. It directly interacts with many members of the SWI/SNF chromatin remodeling complex, including osa, brahma, HDAC3, and moira, along with cytoplasmic members of canonical Wnt signaling pathways CtBP and dishevelled. The SWI/SNF complex (Trithorax group proteins) is thought to be regulated by groucho, which typically represses Wnt target gene expression in the absence of nuclear localized beta-catenin. ebi is homologous to human TBL1X.

This protein is predicted to positively regulate Wnt, N, and EGFR signaling, so we expected that *ebi* RNAi should result in decreased Wnt signaling, paralleling Wnt ligand knockdowns (loss of red patches, etc.)

***ebi expression***

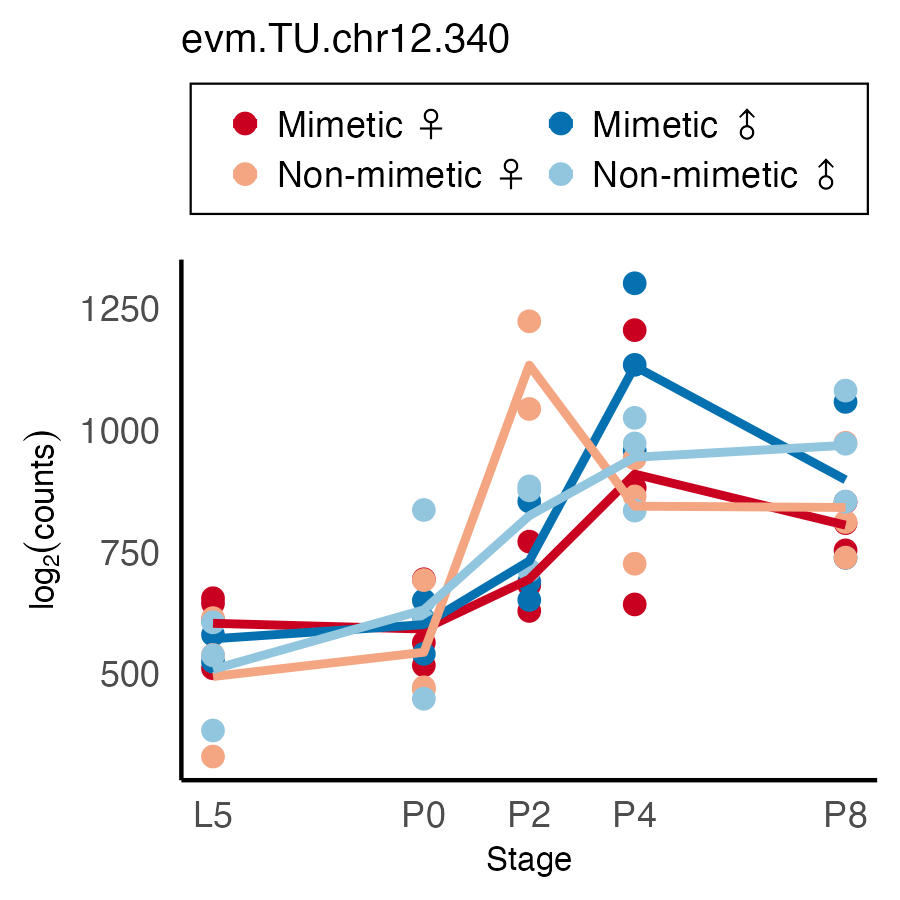

**Phenotype Summary**

RNAi has subtle effects in most cases, but the results suggest that *ebi* normally strongly represses opal scale formation in Cu2-Cu1 cell in mimetic females. Weaker effect on the location of Cu2-Cu1 red patch in mimetic females. Normally represses orange scale formation in proximal melanic regions (especially the DC) in non-mimetic color patterns. Some other small effects on frequency of red scales in A-Cu2 red patch and other marginal crescents, where loss of *ebi* results in more red scales.

**Mimetic Females**

**MF1:**

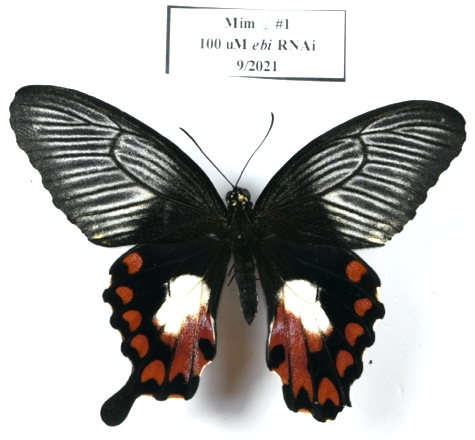

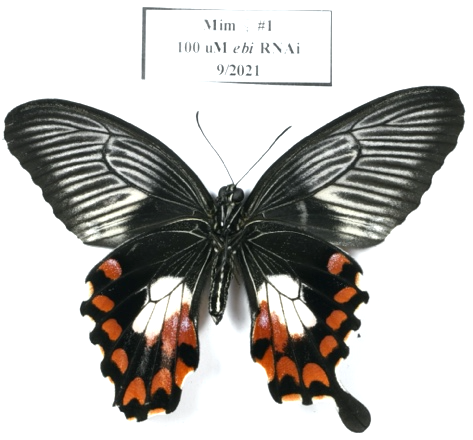

Dorsal:

- Significant increase in opal scales along Cu2 vein in red regions
- Slight distal expansion of white into red region in Cu2-Cu1
- Larger white patch in A-Cu2 cell
- Marginal crescents in Cu2-Cu1, Cu1-M3 much redder.

Ventral:

- Cu2-Cu1 red patch shifted posteriorly (larger white patch)

**MF2:**

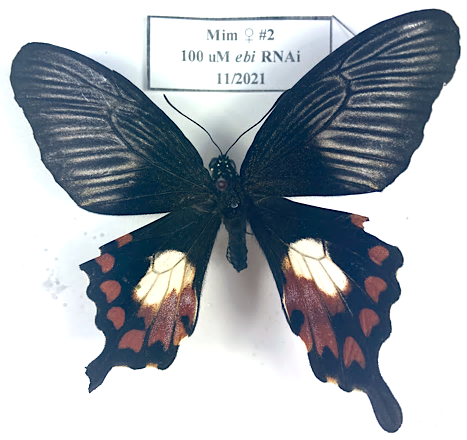

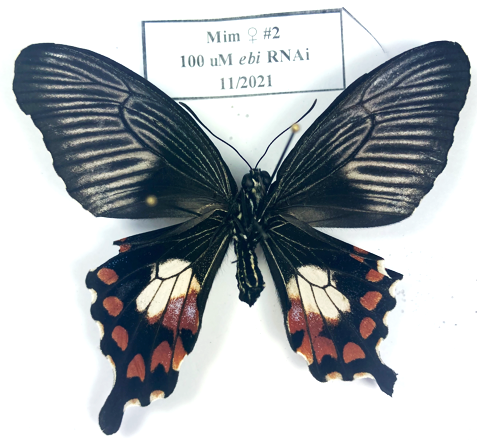

Dorsal:

- Significant increase in opal scales along Cu2 vein in red regions
- Slight distal expansion of white into red region in Cu2-Cu1
- Larger white patch in A-Cu2 cell
- Melanic spot decreased, AT more orange

Ventral:

- Significant increase in red scales along Cu2 vein connecting SMCs and red patches in A-Cu2, Cu2-Cu1 cells
- Significantly FEWER opal scales in red regions in same cells

**MF3:**

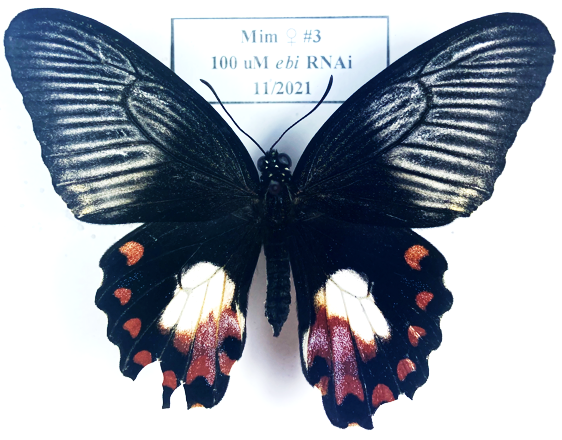

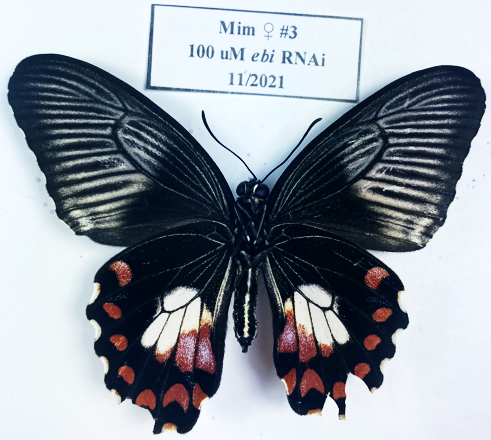

Dorsal:

- Cu2, Cu1 veins significantly more orange scales
- White patch in A-Cu2, Cu2-Cu1 larger (expanded A and P, especially along veins)

Ventral:

- Significantly FEWER opal scales in A-Cu2, Cu2-Cu1 red regions

**Mimetic Males**

**MM1:**

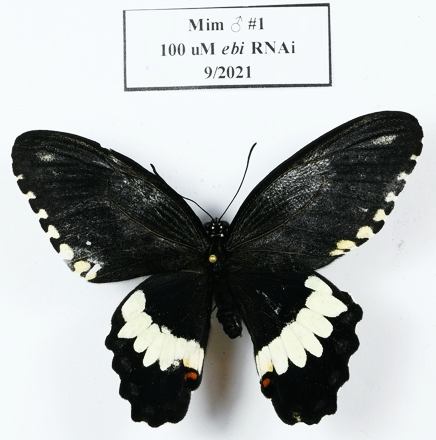

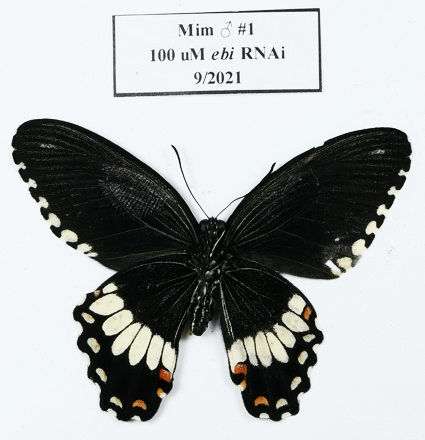

Dorsal:

- Probably mostly damage from injection. Perhaps slight incursion of white into DC.

Ventral:

- Increased frequency of orange cells in distal 1/3^rd^ of DC
- Slightly more red scales trailing from A-Cu2 red patch

**MM2:**

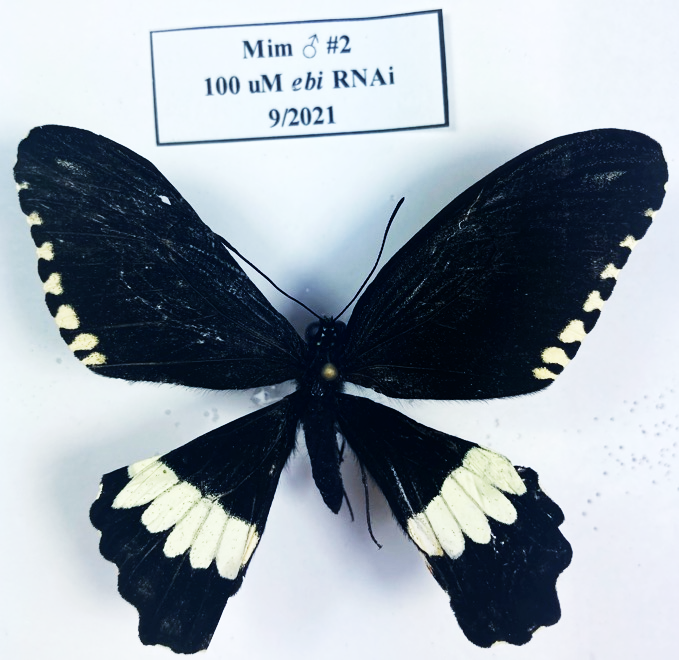

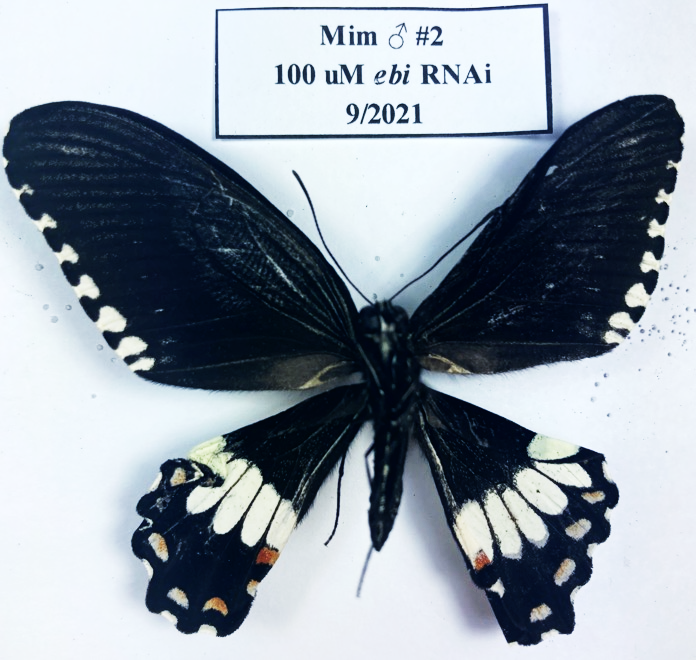

Dorsal:

- Crumpled wings, so hard to tell
- Slightly larger A-Cu2 red patch

Ventral:

- Many fewer red scales in marginal crescents

**MM3:**

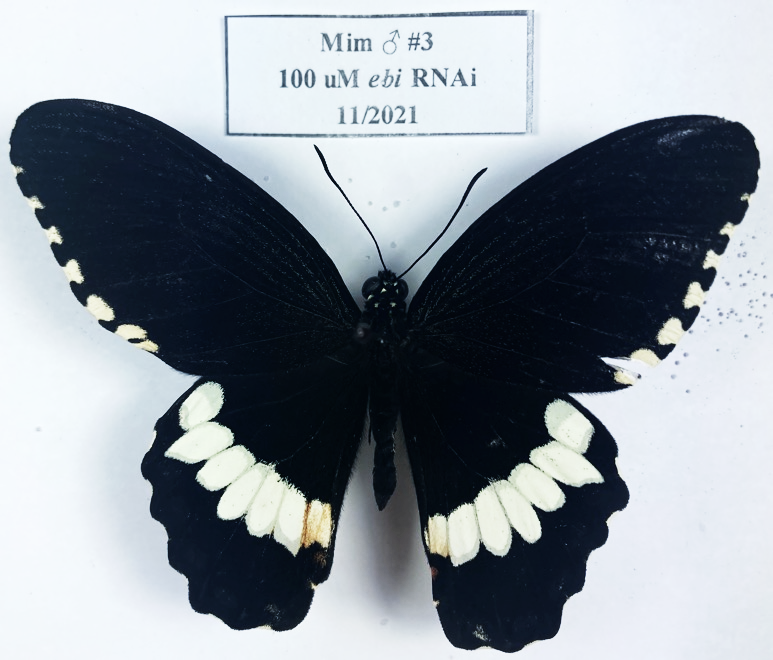

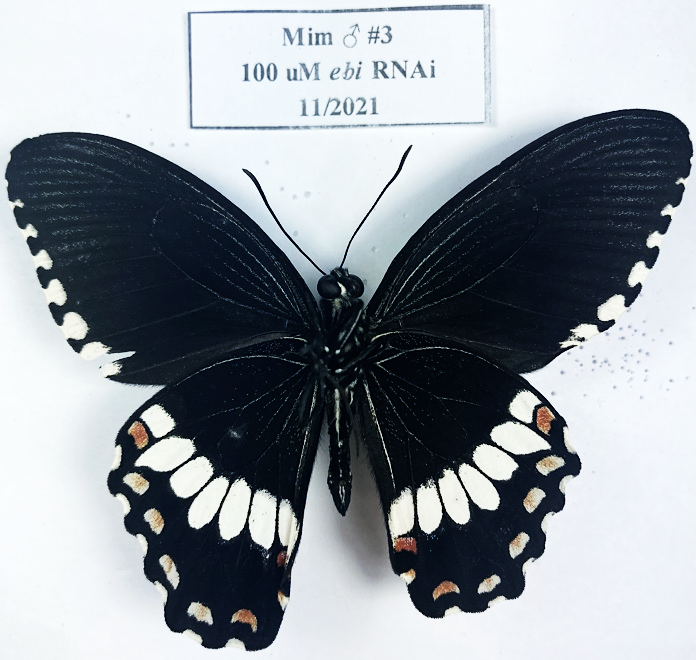

Dorsal:

- Significantly more orange scales in A-Cu2 white patch
- Significantly fewer red scales in A-Cu2 red patch
- Anterior edge of white patches in Cu2-Cu1, Cu1-M3 fuzzy (mixed pale/melanic scales)

Ventral:

- Anterior edge of white patches in Cu2-Cu1, Cu1-M3 fuzzy (mixed pale/melanic scales)

**Non-mimetic Females**

NONE YET

**Non-mimetic Males**

**NM1:**

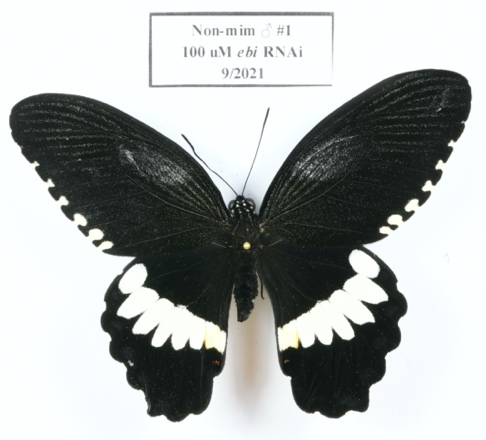

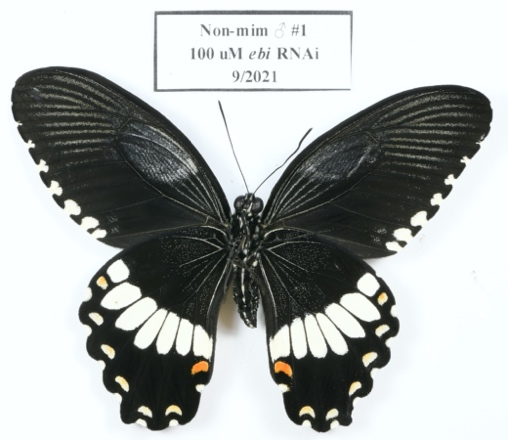

Dorsal:

- NND

Ventral:

- Slight proximal expansion of white patches in cells 2-5.
- Increased frequency of orange scales in posterior half of DC
- Few more red scales trailing from lateral edge of A-Cu2 red patch

**NM2:**

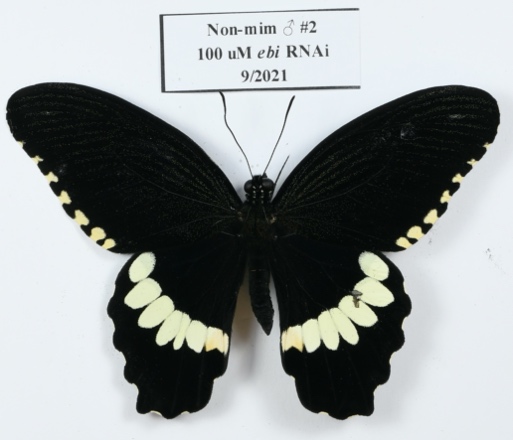

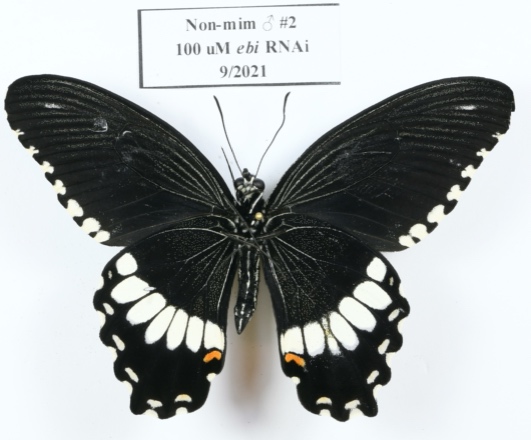

Dorsal:

- NND (*Some damage from injection cased the Cu1-M3 cell to pinch in the white patch.)

Ventral:

- Increased frequency of orange scales in injected region, particularly noticeable in posterior half of DC and just distal to the Cu2-Cu1 white patch.
- Slightly more red scales trailing from lateral edge of A-Cu2 red patch

**NM3:**

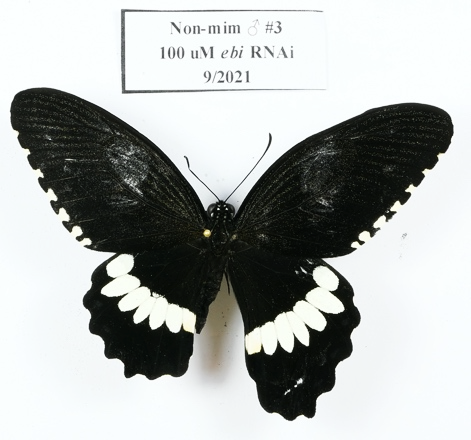

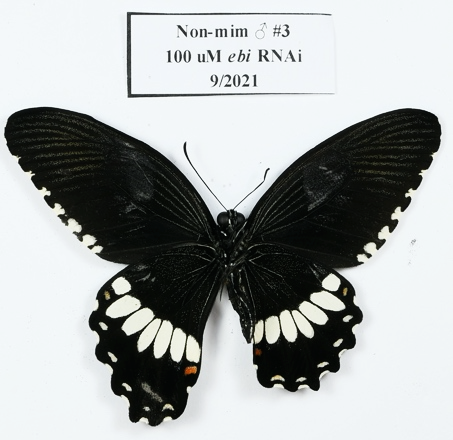

Dorsal:

- Slight anterior expansion of white in DC (could be damage from injection, though).

Ventral:

- Increased frequency of orange cells in distal half of DC

**NM4:**

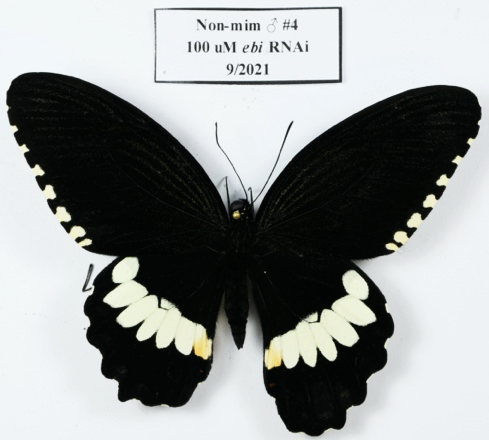

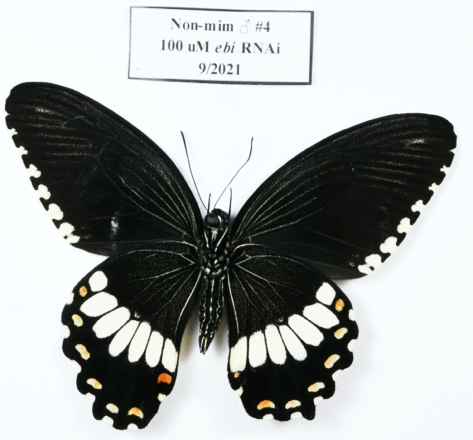

Dorsal:

- Slightly smaller A-Cu2 red patch

Ventral:

- Very small A-Cu2 red patch
- Many more orange scales in DC and into Cu1-M3

***Pygopus (pygo)***

*Pygopus* is a transcriptional co-activator that binds directly to *β-catenin* and His3 in the nucleus to promote *wg*-stimulated transcription of target genes. *Pygo* is part of the main effector complex for Wnt signaling, the TCF/LEF complex. Pygo is thought to help anchor β-catenin in the nucleus, promoting Wnt-mediated transcriptional regulation.

***pygo expression***

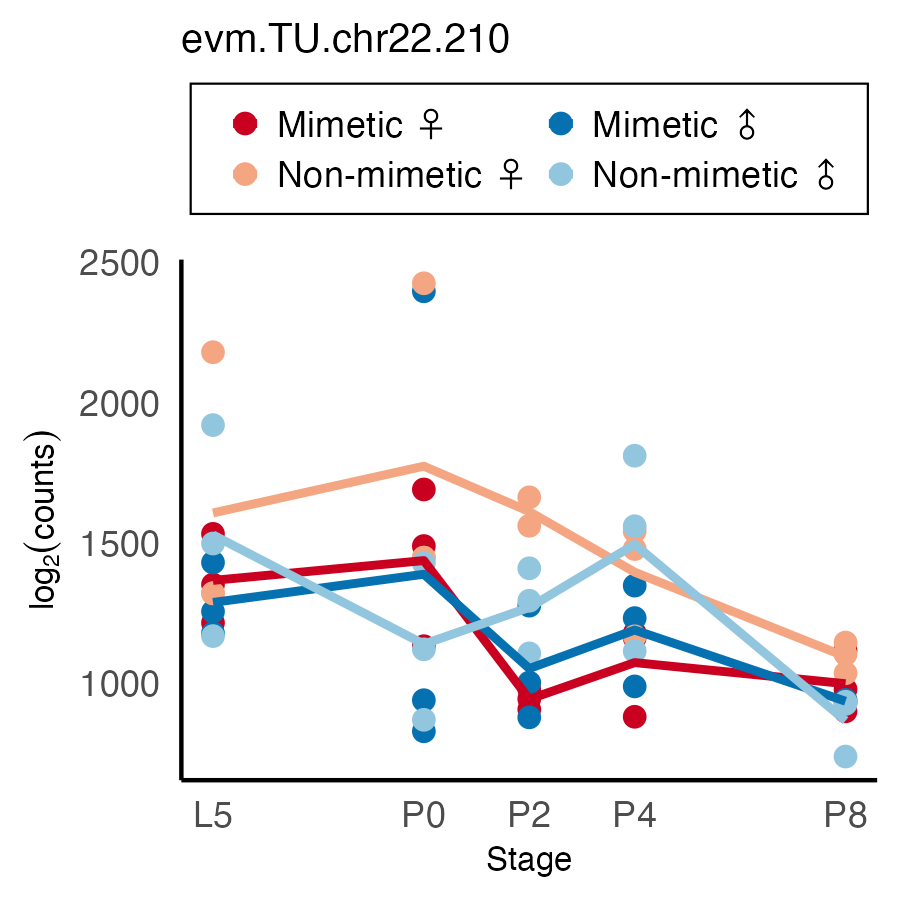

**Phenotype Summary**

RNAi in non-mimetic butterflies results in red scales connecting the A-Cu2 red patch and SMC. Furthermore, *pygo* loss results in slight posterior shift of the anterior edges of the white patches. The posterior boundaries appear to be unchanged. Other slight effects on SMC size and redness. Mimetic female phenotypes resemble those of GSK-3B knockdowns.

**Mimetic Females**

**MF1:**

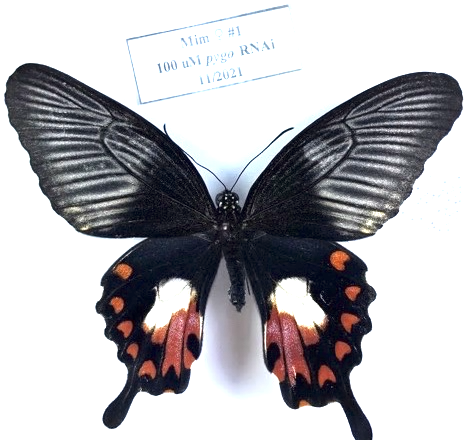

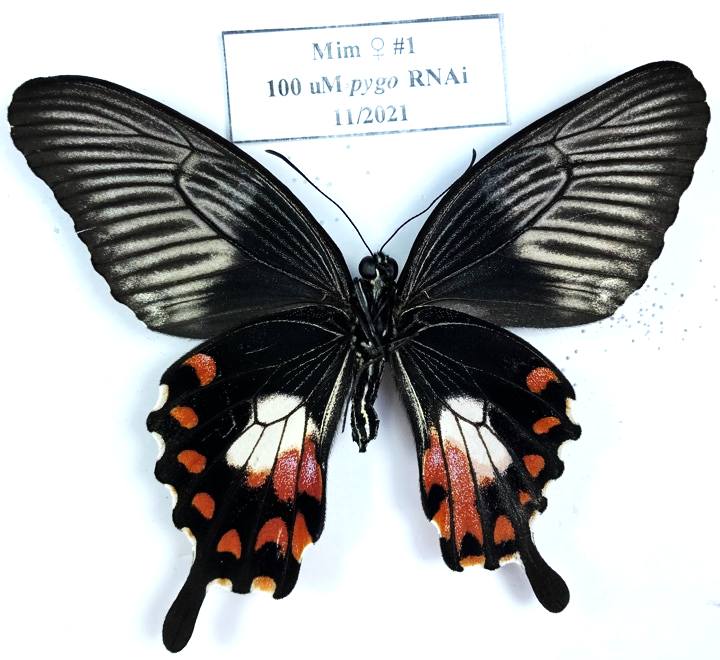

Dorsal:

- Slightly more red scales connecting SMC and red patch in Cu2-Cu1 cell
- Significantly fewer opal scales in A-Cu2, Cu2-Cu1 cells

Ventral:

- Loss of white patch in A-Cu2
- Red scales connecting red patch and AT in A-Cu2 cell
- Red scales fully connecting SMC and red patch in Cu2-Cu1 cell, SMC edges “trail” posteriorly. Strangely shaped marginal patch. Anterior edge of red patch moved anteriorly.
- Cu1-M3 white patch extended posteriorly

**MF2:**

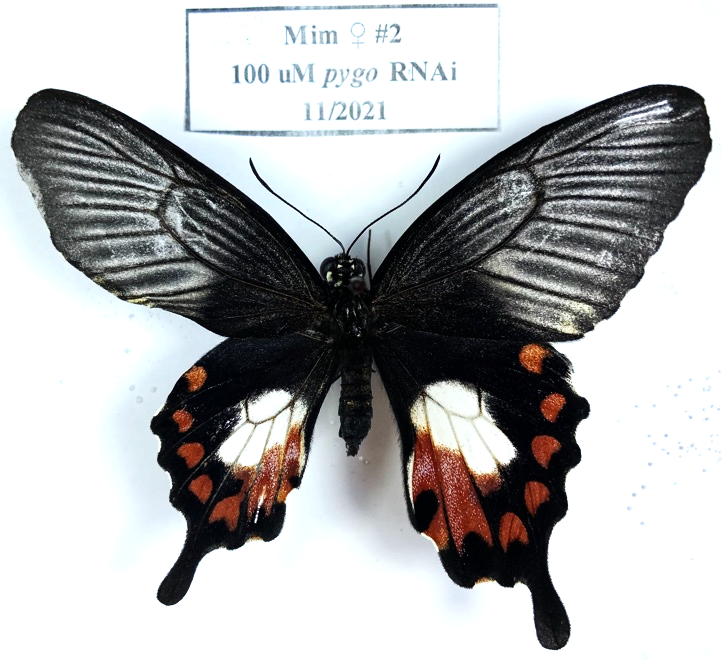

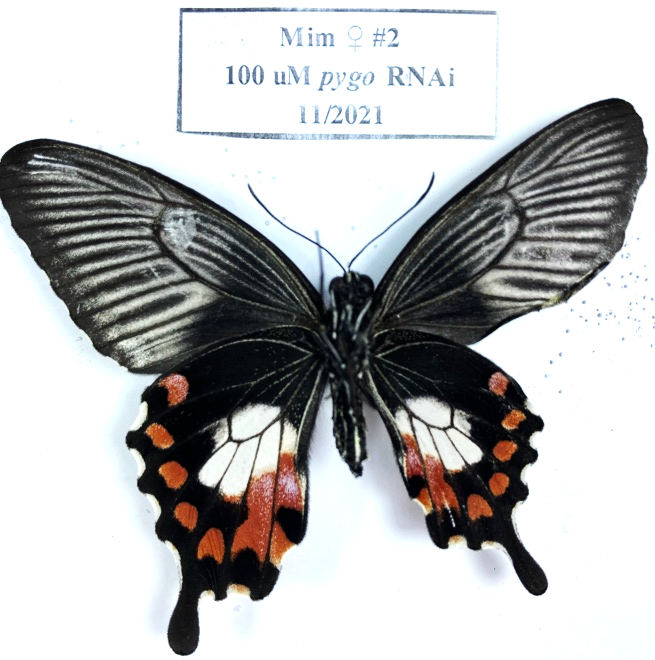

Dorsal:

- Cu1-M3 red patch connected to SMC along the Cu1 vein.

Ventral:

- NND

**Mimetic Males**

**MM1:**

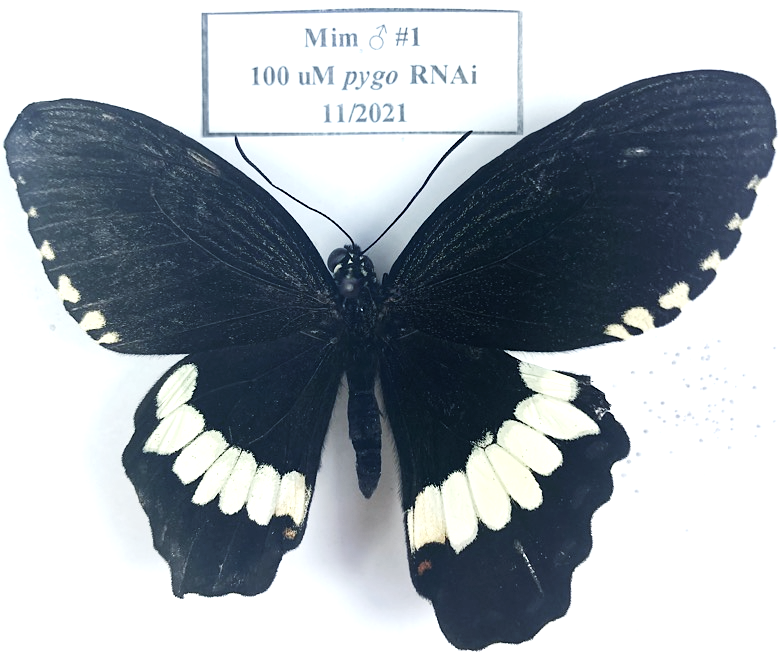

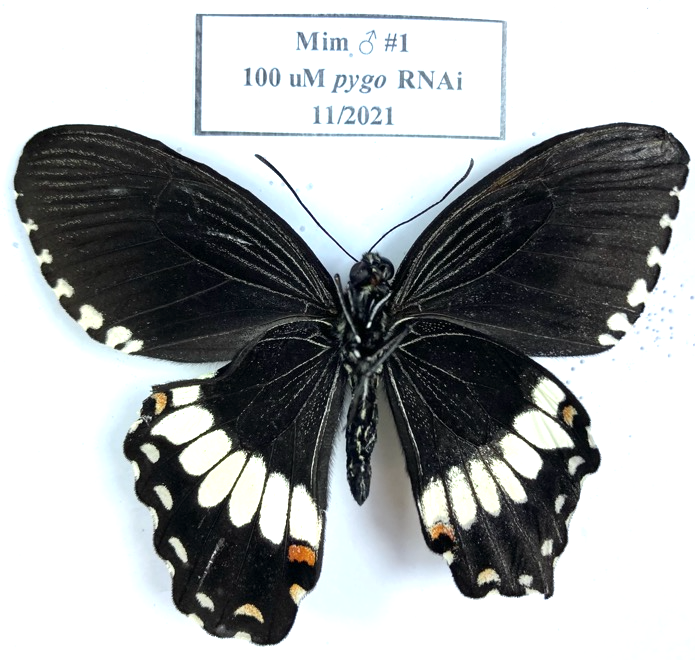

Dorsal:

- Fewer orange scales in A-Cu2 pale patch
- A-Cu2 red patch smaller and diffuse with melanic scales
- No pale scales in DC

Ventral:

- Melanic scales infiltrate edges of pale patches in first 5 cells
- Red patch in A-Cu1 cell infiltrated with pale scales

**MM2:**

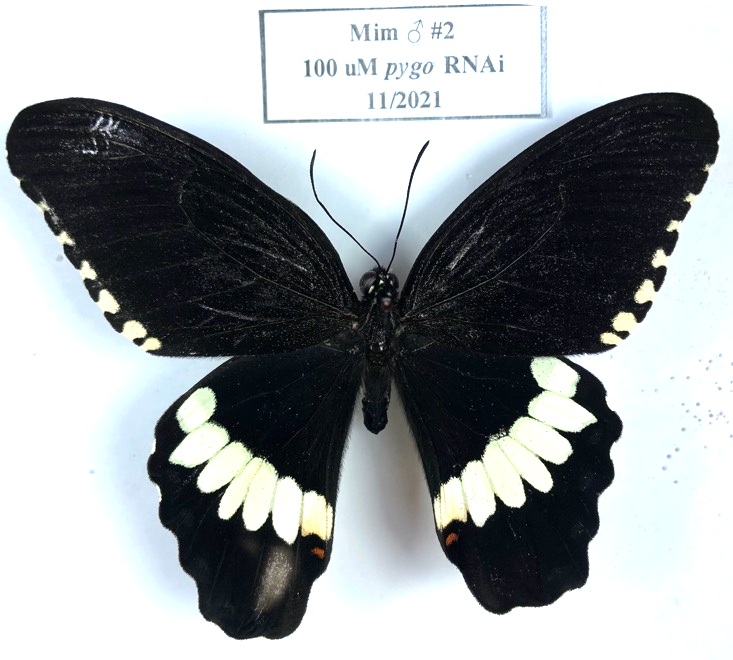

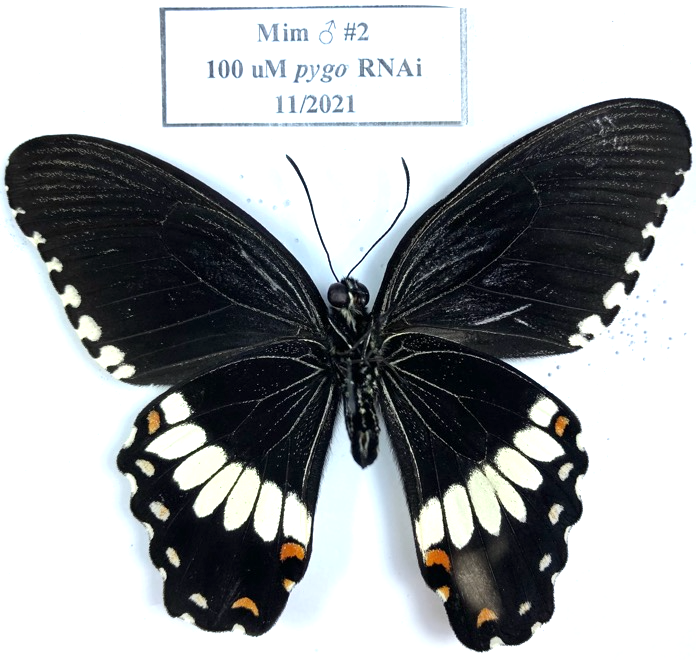

Dorsal:

- NND, except gray patch (far from injection site) that appears on both D and V… not sure, never seen it before. Assume it’s damage.

Ventral:

- NND

**MM3:**

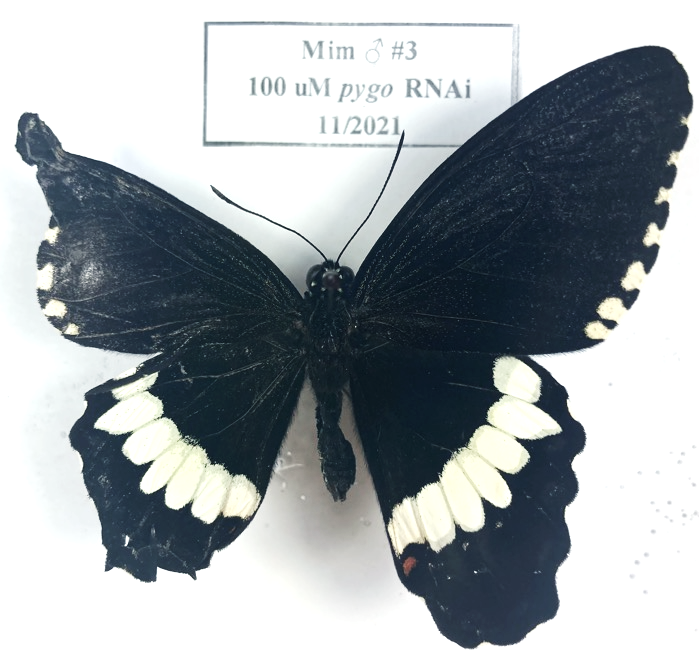

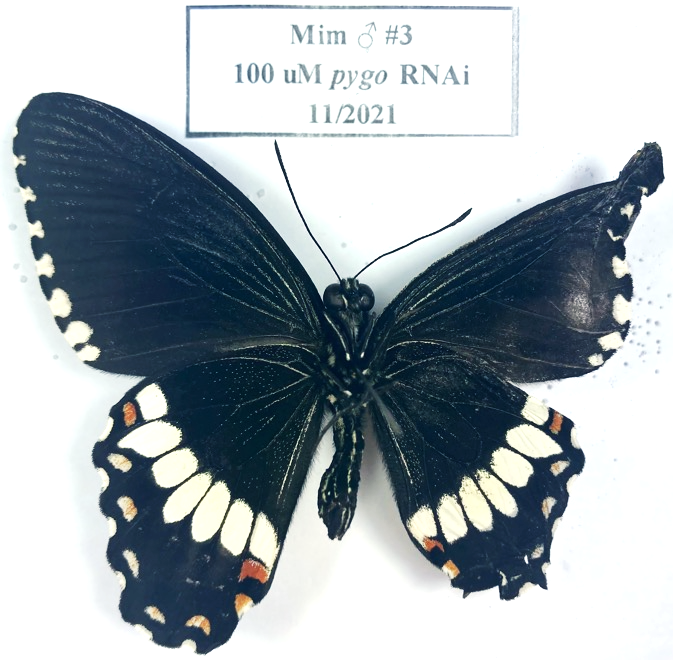

Dorsal:

- Lots of damage during eclosion, but appears to be NND

Ventral:

- NND

**Non-mimetic Females**

**NF1:**

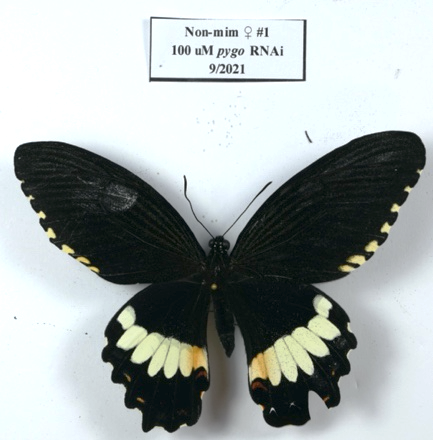

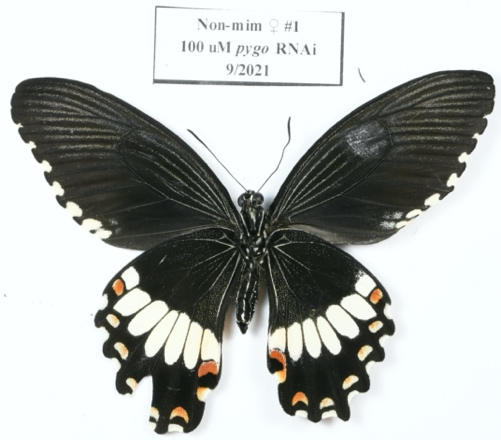

Dorsal:

- A-Cu2 rusty patch much redder – more red scales

Ventral:

- Anterior edge of white patches in medial 4 cells are shifted posteriorly
- Slight increase in orange scales in distal DC
- A-Cu2 red patch and AT connected by rows of red and opal scales
- Increased frequency of opal scales in Cu2-Cu1 posterior to white patch

**NF2:**

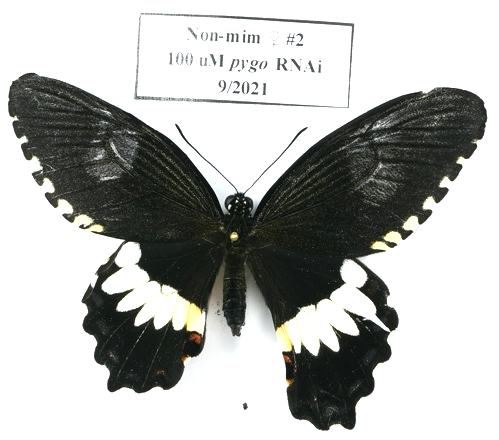

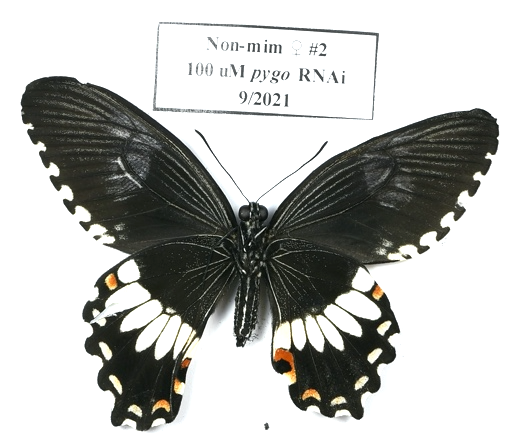

Dorsal:

- Slightly larger SMC in Cu2-Cu1

Ventral:

- Strong connection between A-Cu2 red patch and AT, filled in with red scales
- Slight posterior shift of the anterior edge of white patches in medial 5 cells

**NF3:**

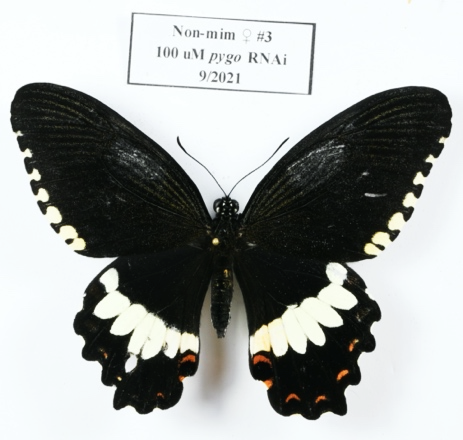

Dorsal:

- Posterior shift of the anterior edge of white patch in medial 4 cells
- Reduction or loss of SMCs in cells 2-5

Ventral:

- Strong connection between A-Cu2 red patch and AT, filled with red scales
- Cu1-M3, M3-M2 SMCs small, all white (could be partially from injection damage)
- Slight posterior shift for anterior edges of Cu1-M3, M3-M2 white patches

**Non-mimetic Males**

**NM1:**

Dorsal:

- A-Cu2 red patch much larger, trails to AT
- Anterior edges of white patches in medial 4 cells shifted slightly posteriorly

Ventral:

- Anterior edges of white patches in medial 5 cells shifted slightly posteriorly
- Red scales connecting A-Cu2 red patch and AT. Some opal scales.

***Engrailed (en)***

*Engrailed* is a homeodomain-containing transcription factor essential for posterior compartment identity, particularly well studied in *Drosophila* wing development. En is the main transcription factor effector activated Hh signaling, and bothpositively regulates Hedgehog signaling and negatively regulates Hh targets such as *cubitus interruptus, patched,* and *decapentaplegic*. *En* and its paralog *invected* are implicated in the *Papilio dardanus* mimicry switch.

***en expression***

**Phenotype Summary**

Phenotypic effects of *en* differ between mimetic females and all other groups in several ways. First *en* RNAi results in the mimetic female white patch shrinking and the majority of white scales developing as opal and gray scales. Mimetic females also lost all red scales except the A-Cu2 red patch and AT, which comes to resemble the non-mimetic female color pattern in this cell. Regions that are normally white or red become melanic. Furthermore, loss of *en* results in a significant increase in opal scales in the medial two cells, so that those regions appear white in the pictures below. Mimetic females still do not develop pale yellow scales, suggesting that *en* is not restricting pale scale fate. Finally, the effects are similar on both dorsal and ventral color patterns.

In contrast, *en* RNAi in butterflies with non-mimetic color patterns causes different effects on the dorsal and ventral surfaces. Dorsally, *en* RNAi causes the medial 2 – 3 pale yellow patches to shift posteriorly and trail off posteriorly into the melanic regions. Ventrally, *en* RNAi causes the entire pattern to shift posteriorly, frequently resulting in loss of SMCs or at least merging of SMCs with marginal crescents. Again, scale colors do not appear to change. I observed no effects on opal scale presence or frequency in non-mimetic patterned bugs.

We should note that most of the A-P/D-V axis information has already been laid down by 5^th^ instar and that we should be mainly picking up the effects of *en* from pupation onwards (see expression above). Low expression in non-mimetic patterned bugs is also consistent with the generally weaker effects of *en* RNAi in these individuals.

**Mimetic Females**

**MF1:**

Dorsal:

- DC: anterior edge moved posteriorly (overall appears to have “shrunk”)
- A-Cu2: loss of almost all white and orange scales, anterior boundary of red patch shifted slightly posterior. Large increase in frequency of opal scales.
- Cu2-Cu1: White patch shrunken, infused with some melanic scales. Anterior boundary of red patch shifted slightly posterior, but otherwise unchanged. Large increase in frequency of opal scales.
- Cu1-M3: White patch shrunken particularly at posterior edge, infused with some melanic scales. Red patch shrunken, square.
- M3-M2: White patch shrunken, particularly away from posterior boundary. Loss of all red scales.

Ventral:

- Overall:
  - appears that the proximal pattern (melanic, speckled with orange scales) extends distally past the border of the DC.
  - Melanic regions near injection site appear gray – perhaps ground scales are changed color?
  - SMCs unchanged, perhaps because injection was localized (i.e. I don’t see gray scales near the SMCs).
- DC: complete loss of white scales, only melanic with orange speckled.
- A-Cu2: Complete loss of white scales.
  - Red patch almost completely converted to opal scales.
  - Red patch and AT strongly connected by opal (proximal) and red (distal) scales.
  - Melanic / speckled orange extends distally to fill in normally white regions.
- Cu2-Cu1: Almost complete loss of white.
  - No red scales remain.
  - Large number of opal scales in region that is normally red. Roughly even with number of melanic scales in this region.
- Cu1-M3: White patch shrunken, replaced with melanic scales.
- M3-M2: White patch shrunken, replaced with melanic scales.
- M2-M1: Complete loss of white scales.

**MF2:**

Dorsal:

- Overall:
  - Melanic regions 🡪 “gray” (think ground scales changed/lost color)
  - White patch appears shrunken
  - Serious increase in opal scale frequency
- A-Cu2:
  - Complete loss of white scales, most orange scales from proximal region.
  - Red patch lost except for the top of the hook, as in non-mimetic females.
  - Increased frequency of opal scales, particularly in red regions and connection between red patch and AT
- Cu2-Cu1:
  - Many more opal scales
  - White patch shrunken
  - Complete loss of red scales
  - Reduced SMC size
- Cu1-M3:
  - Many more opal scales
  - White patch shrunken
  - Complete loss of red scales
  - Reduced SMC size
- M3-M2:
  - Many more opal scales
  - White patch shrunken
  - Complete loss of red scales
  - Reduced SMC size

Ventral:

- Overall:
  - White patch shrunk
  - Complete loss of red patches in all cells except A-Cu2, but this is differently shaped and looks like non-mimetic ventral pattern
  - Significant increase in opal scales
- DC:
  - White patch shrunk, anterior edge shifted posteriorly
- A-Cu2:
  - Complete loss of white
  - Red patch shaped like non-mimetic pattern, but patch and AT joined by red scales.
  - Significant increase in opal scales throughout proximal 2/3 of the red region
- Cu2-Cu1:
  - Complete loss of red scales
  - Proximal edge of white patch shifted posteriorly, patch overall smaller
  - Posterior edge of white patch fuzzy, mixed white and melanic scales
- Cu1-M3:
  - White patch shrunk, posterior edge shifted anteriorly
  - Complete loss of red scales
  - Posterior edge of white patch fuzzy, mixed white and melanic scales

**Mimetic Males**

**MM1:**

Hard to tell because crumpled.

Dorsal:

- NND

Ventral:

- Anterior border of white patches in medial 5 cells shifted posteriorly. Posterior boundary unchanged.
- New melanic portions are gray, not background black.
- Melanic/orange speckle pattern extends posteriorly in DC and medial 4 cells

**MM2:**

Dorsal:

- Slightly more red scales in A-Cu2 AT/red patch
- Anterior boundary of A-Cu2 white patch shifted posteriorly
- Posterior boundaries of medial 4 cells a little fuzzy (mixed melanic/pale)

Ventral:

- Overall:
  - White patches in medial 5 cells shifted posteriorly.
  - **INTERESTINGLY, the newly white regions do not fluoresce under UV, but the regions that are normally pale still do! I am not sure if this is because the fluorescence is coming from the dorsal scales, though, and that it appears that there is less fluorescence in the posterior regions because there are no fluorescent scales on the dorsal side there. The scales in the two regions have the same morphology and apparent color under the dissecting scope. Maybe just bleed through.
  - Opal scales still show up in A-Cu2 region that corresponds to the red patch.
  - SMCs and marginal crescents merged in medial 5 cells, mimetic white
  - Almost all red lost from marginal crescents
  - A-Cu2 red patch and AT turned white, but sprinkled with opal scales like normal. Melanic spot maintained.

**MM3:**

Dorsal:

- White patches in medial 4 cells shifted posteriorly, with extension along Cu2 vein of pale yellow scales.
- Expanded AT with orange and white scales
- A-Cu2 red patch covered by pale scales

Ventral:

- White patches in medial 5 cells shifted strongly posterior
- Cu2-Cu1, Cu1-M3 SMCs significantly reduced.
- No red scales in SMCs in cells 2-6
- AT extended anterior with orange and white scales

**MM4:**

Dorsal:

- Slightly more orange scales connecting AT and red patch in A-Cu2

Ventral:

- Anterior boundaries of white patches in medial cells 2 – 5 shifted posteriorly.
- Melanic/speckled orange region extends to distal DC
- SMC and marginal crescent in Cu2-Cu1 beginning to merge

**Non-mimetic Females**

**NF1:**

Dorsal:

- A-Cu2 red patch smaller, more orange scales
- A-Cu2 pale patch has fewer orange scales

Ventral:

- Anterior boundaries of medial cells 2-4 shifted posteriorly, fuzzy due to increased melanic scales. Generally smaller.
- Opal scales throughout distal melanic region in Cu2-Cu1, Cu1-M3
- Slightly more orange scales in distal DC

**NF2:**

Dorsal:

- Anterior boundaries of pale patches in medial 3 cells shifted posterior.
- Complete loss of orange scales (and fluorescence) in Cu2-Cu1 patch
- Red significantly reduced (could just be because pattern is shifted posteriorly, so those SMCs are pushed off the wing/squished)
- Posterior boundaries of those pale patches are fuzzy – infused with many melanic scales

Ventral:

- Slight posterior shift of anterior boundaries of white patches in medial 2-3 cells
- Loss of orange scales in Cu2-Cu1 cell

**Non-mimetic Males**

**NM2a:**

Dorsal:

- NND

Ventral:

- White patches strongly shifted posterior in medial 5 cells
- A-Cu2 white patch overrides red patch, white scales connect red patch and AT
- Corresponding expansion of melanic/orange speckle region in Cu2-Cu1 and DC (slight)
- Complete loss of orange scales in all SMCs
- SCMs reduced and diffuse in Cu2-Cu1 and Cu1-M3 cells

**NM2b:**

Dorsal:

- Anterior boundaries of pale patches in medial 3 cells shifted posteriorly
- Loss of orange scales in medial 2 cells
- Trailing posterior edges of pale patches in medial 2 cells

Ventral:

- Anterior boundaries of medial 3 cells slightly shifted posterior and fuzzy

**NM3a:**

Dorsal:

- NND

Ventral:

- Maybe weak shift of pale patches to the posterior in medial cells 2-4

**NM3b:**

Dorsal:

- Anterior boundaries of pale patches in medial 3 cells shifted posteriorly, fuzzy due to mix of melanic and pale scales
- Posterior boundaries of these same patches fuzzy, trailing
- Loss of orange scales in Cu2-Cu1 cell
- Loss of pale scales in DC (probably due to posterior shift of patch)

Ventral:

- Cu1-M3 pale patch slight incursion of melanic scales in medial-anterior corner

**NM4:**

Dorsal:

- Anterior boundaries of pale patches in medial cells 1-5 shifted posteriorly
- Pale patches extended posteriorly in medial 4 cells, trailing
- Loss of orange scales in Cu2-Cu1 pale patch

Ventral:

- Very slight posterior shift of pale anterior boundary in Cu1-M3 cell

***Invected***

*invected* is a homeodomain-containing transcription factor essential for posterior compartment identity, particularly well studied in *Drosophila* wing development. Inv is thought to positively regulate Hedgehog signaling and negatively regulate Hh targets such as *cubitus interruptus, patched,* and *decapentaplegic*. *Inv* and its paralog *engrailed* are implicated in the *Papilio dardanus* mimicry switch.

***inv expression***

**Phenotype Summary**

Few *invected* RNAi individuals survived to emergence, so our data is limited. However, it is clear that *inv* has significantly different effects than *en*. *inv* RNAi caused some mimetic color pattern elements to revert to a non-mimetic like form, particularly the pattern in the A-Cu2 cell.

**Mimetic Females**

**MF1 (4/2022 – 100 uM)**

Dorsal:

- Posterior expansion of central white patches over red patches
- Loss of marginal spots in Cu2-Cu1 and Cu1-M3 cells
- Slight distal extension of M3-M2 white/red patch

Ventral:

- Distal expansion of white patches in Cu2-Cu1, Cu1-M3, and M3-M2 wing cells.
- Reduced sized of AT and Cu2-Cu1 and Cu1-M3 SMCs
- Distal shift of the anterior edge of the A-Cu2 red patch and reduced frequency of red scales, causing it to resemble the non-mimetic female pattern in this region.

***Hedgehog (hh)***

**Phenotype Summary**

Few *hh* RNAi individuals emerged, making it difficult to draw any conclusions. Effects were weak.

**Mimetic Females**

**MF1**

Dorsal:

- Distal extension of white patches over red in wing cells 2-4, somewhat resembling *inv* RNAi
- Slight distal extension of red patches in those same cells

Ventral

- Increased red scales connecting AT and red patch in A-Cu2 cell
- Slight distal extension of white patches in medial four wing cells

**Mimetic Males**

**MM1**

Dorsal:

- NND

Ventral:

- Slight posterior shift of pale patches in wing cells 2-4, causing SMCs to merge with MCs, reminiscent of *en* RNAi phenotype

***Decapentaplegic (dpp)***

*Dpp* is one of three ligands in the BMP branch of the TGF-B signaling pathway and serves critical roles specifying the proximodistal axis in the developing *Drosophila* wing.

***dpp expression***

**Phenotype Summary**

**Mimetic Females**

**MF1**

Dorsal:

- Increased red scales connecting AT and A-Cu2 red patch.
- Increase in gray scales in central white patch
- Slight distal extension of central red patches

Ventral:

- Significant increase in red scales in A-Cu2, Cu2-Cu1 cells
- Significant increase in gray scale frequencies in the central white patch

**Mimetic Males**

**MM1**

Dorsal:

- NND

Ventral:

- NND

**MM2**

Dorsal:

- NND

Ventral:

- NND

**MM3**

Some damage from poor emergence, so cannot be certain of phenotypes.

Dorsal:

- NND

Ventral:

- NND

**MM4**

Dorsal:

- NND

Ventral:

- NND

**Non-mimetic Females**

**NF1**

Dorsal:

- Reduced red SMCs
- Slight distal shift of pale band in medial 4 wing cells

Ventral:

- Loss of Cu2-Cu1SMP
- Slight increase in gray scales in Cu2-Cu1 pale patch

**Non-mimetic Males**

**NM1**

Dorsal:

- Loss of pale scales in DC
- Reduced frequency of orange scales in A-Cu2 pale patch

Ventral:

- Narrow stripe of red scales connecting AT to A-Cu2 red patch
- Increased red/orange scales in Cu2-Cu1 SMC

**NM2**

Dorsal:

- Anterior edge is folded over, not different pattern
- NND

Ventral:

- NND

***Doublesex exon 3***

**Mimetic Females**

**MF2**

Dorsal: A little difficult to tell because wing was distorted during emergence.

- Conversion of white scales to pale scales in the central region
- Loss of central red patches
- Conversion of A-Cu2 red and white pattern to the non-mimetic-like pattern.
- Loss of red scales in SMC, converting to non-mimetic like color

Ventral:

- Loss of central red patches
- Loss of red scales in SMCs
- Central white patch extended distally, but mostly due to problems during emergences

**MF4**

Shown in main figure

**Mimetic Males**

**MM1**

Dorsal:

- NND

Ventral:

- A few extra red scales connecting the AT and A-Cu2 red patch

**MM2**

Dorsal:

- NND

Ventral:

- NND

***Doublesex exon 5***

**Mimetic Females**

**MF1**

Dorsal:

- Slight distal extension of Cu2-Cu1 red patch

Ventral:

- Distal shift of proximal edge of white patches in medial two wing cells and DC
- Distal expansion of distal edges of white patches in wing cells 2-5, overriding red patches
- Distal shift of proximal edge of A-Cu2 red patch

**MF2**

Dorsal:

- Increased red scales along Cu2 vein, extending red patches
- Distal expansion of M2-M1 white patch

Ventral:

- Distal shift of proximal edge of white patches in medial two wing cells and DC
- Gain of white scales in M1-Sc+Rs cell where non-mimetic band appears
- Distal expansion of distal edges of white patches in wing cells 2-5, overriding red patches
- Distal shift of proximal edge of A-Cu2 red patch

**Mimetic Males**

**MM1**

Dorsal:

- NND

Ventral:

- Slight distal expansion of pale patches in wing cells 3 and 4

**MM2**

Dorsal:

- NND

Ventral:

- Red scales connecting AT and A-Cu2 red patch

***Doublesex exon 5 (non-mimetic)***

**Non-mimetic Females**

**NF1**

Dorsal:

- Potentially smaller SMCs, although could be from damage during emergence

Ventral:

- Smaller SMCs
- Slight increase in pale scales in DC

**Non-mimetic Males**

**NM1**

Dorsal:

- Large increase in the orange/pale scales in A-Cu2 cell, much more like non-mimetic female
- Appearance of small red patch in A-Cu2

Ventral:

- Some damage from emergence and injection itself, but certainly fuzzy boundaries to central pale patches, infused with melanic scales

**NM3**

Dorsal:

- Large increase in the orange/pale scales in A-Cu2 cell, much more like non-mimetic female
- Appearance of small red patch in A-Cu2

Ventral:

- Red scales connecting AT and A-Cu2 red patch
- Pale scales appeared in the DC
- Cu2-Cu1 SMC more red scales

**NM3b**

Dorsal:

- Large increase in the orange/pale scales in A-Cu2 cell, much more like non-mimetic female

Ventral:

- Cu1-M3 SMC slightly shifted distally
- Slight increase in white scales in DC

***Tyrosine Hydroxylase* (*TH*)**

Very early injections to test for the efficacy of the RNAi. Demonstrates that the DsiRNAs reside in the tissue until at least 13 days after pupation, the first time when TH is turned on (Fujiwara paper describing development of color pattern).
